# Tau Secretion and Propagation Is Regulated by p300/CBP via Autophagy-Lysosomal Pathway in Tauopathy

**DOI:** 10.1101/418640

**Authors:** Xu Chen, Yaqiao Li, Chao Wang, Yinyan Tang, Sue-Ann Mok, Richard M. Tsai, Julio C. Rojas, Anna Karydas, Bruce L. Miller, Adam L. Boxer, Jason E. Gestwicki, Ana Maria Cuervo, Michelle Arkin, Li Gan

**Affiliations:** Gladstone Institute of Neurological Disease, University of California, San Francisco, CA 94158, USA; Department of Neurology, University of California, San Francisco, CA 94158, USA; Small Molecule Discovery Center, Department of Pharmaceutical Chemistry, University of California, San Francisco, CA 94158, USA; Institute for Neurodegenerative Disease, Department of Pharmaceutical Chemistry, Weill Institute for Neurosciences, University of California, San Francisco, CA 94158, USA; Memory and Aging Center, University of California, San Francisco, CA 94158, USA; Department of Developmental and Molecular Biology, Albert Einstein College of Medicine, Bronx, NY 10461, USA; Institute for Aging Studies, Albert Einstein College of Medicine, Bronx, NY 10461, USA.

## Abstract

The trans-neuronal propagation of tau has been implicated in the progression of tau-mediated neurodegeneration. Tau secretion from neurons is the first step in tau transmission, but little is known about the cellular mechanism. Here, we report that p300/CBP, the lysine acetyltransferase that acetylates tau and regulates its homeostasis and toxicity, serves as a key regulator of tau secretion by inhibiting the autophagy-lysosomal pathway (ALP). Increased p300/CBP activity was associated with impaired function of this pathway in a tau transgenic mouse model. p300/CBP hyperactivation increased tau secretion by blocking autophagic flux. Conversely, inhibiting p300/CBP genetically or pharmacologically promoted autophagic flux, and reduced tau accumulation, tau secretion, and tau propagation in fibril-induced tau spreading models in vitro and in vivo. Our findings show that p300/CBP-induced impairment in the ALP underlies excessive unconventional secretion and pathogenic spread of tau.

## INTRODUCTION

Tauopathies, including Alzheimer’s disease (AD) and frontotemporal lobe degeneration with tau inclusions, are neurodegenerative diseases characterized by neuronal deposition of neurofibrillary tangles composed of the microtubule-associated protein tau. Tauopathy progression is strongly associated with spreading of tau pathology in a stereotypic pattern, which is used to stage the disease (Jucker and Walker, 2013, Braak and Braak, 1991). Cell-to-cell transmission of pathogenic tau species may account for the spreading of tau pathology in the brain (Brettschneider et al., 2015, Iba et al., 2013, Clavaguera et al., 2009). How tau transmits and propagates between neurons is poorly understood.

The intracellular proteostatsis of tau is maintained by the autophagy-lysosomal pathway (ALP) and the ubiquitin-proteasomal system (UPS). Impairment of these degradative systems leads to intracellular tau accumulation, neuronal deficits, and aggregate formation. Tau is also present extracellularly, in the cerebrospinal fluid (CSF) and interstitial fluid of tauopathy brains and in the conditioned medium of neuronal culture. Extracellular tau species can be toxic to the neurons (Diaz-Hernandez et al., 2010, Kaniyappan et al., 2017) and more importantly, can become pathogenic “seeds” to be taken up by surrounding neurons (Wu et al., 2016, Wang et al., 2017b). Rather than being passively released after neuronal death, tau is actively secreted by live neurons (Chai et al., 2012, Karch et al., 2012)—a critical step toward cell-to-cell tau transmission. However, the cellular, molecular, and regulatory mechanisms of tau secretion are largely unknown. It has been proposed that tau, which lacks a signal peptide, might be secreted by unconventional secretion (Chai et al., 2012, Borland and Vilhardt, 2017), a pathway that overlaps with the ALP (Zhang and Schekman, 2013, Ponpuak et al., 2015). Thus, autophagy—a key mechanism for tau degradation—might have an uncharacterized role in tau secretion and propagation.

p300/CBP, a lysine acetyltransferase for numerous nuclear and cytosolic proteins, is a master regulator of Alzheimer’s disease progression (Aubry et al., 2015). Previously, we showed that p300/CBP is the acetyltransferase for tau and that hyperacetylation of tau by increased p300/CBP activity increases tau accumulation (Min et al., 2015, Min et al., 2010). Inhibition of p300/CBP reduces tau accumulation, tau pathology, and cognitive deficits in tau transgenic mice, highlighting a critical pathogenic role of p300/CBP (Min et al., 2015). However, it is not known whether p300/CBP regulates the secretion and propagation of tau during disease progression.

In this study, we investigated the possibility that p300/CBP regulates tau secretion through effects on autophagy. In tauopathy brains, we examined changes in p300/CBP activity and how that is associated with autophagy impairment. Using HEK293T cell reconstitution and neuronal culture, we investigated the relationship between p300/CBP, autophagic flux and tau secretion. We performed a high throughput screen for new p300 inhibitors, and identified a compound that has strong effect on reducing tau secretion. Using this compound and genetic manipulations, we further investigated how p300/CBP inhibition affects tau propagation in both in vitro and in vivo tau spreading models.

## RESULTS

### Hyperactive p300/CBP in Tauopathy Brains Is Associated with Autophagy Impairment

p300/CBP activity is increased in tauP301S transgenic mice (PS19) (Min et al., 2015), a mouse model of tauopathy (Yoshiyama et al., 2007). Using acetylated histone H3 (acH3K18)—a substrate of p300/CBP (Jin et al., 2011)—as a marker for p300/CBP activity, we found that acH3K18 is completely absent in p300/CBP double-knockout primary neurons (Figure S1A, S1C). To assess p300/CBP activity in human tauopathy brains, we measured acH3K18 levels in CSF samples from patients with AD and progressive supranuclear palsy (PSP). In both groups, the levels were significantly higher than in healthy controls, as shown by ELISA (Figure 1A), indicating aberrantly increased p300/CBP activity in the diseased brains.

**Figure 1.**
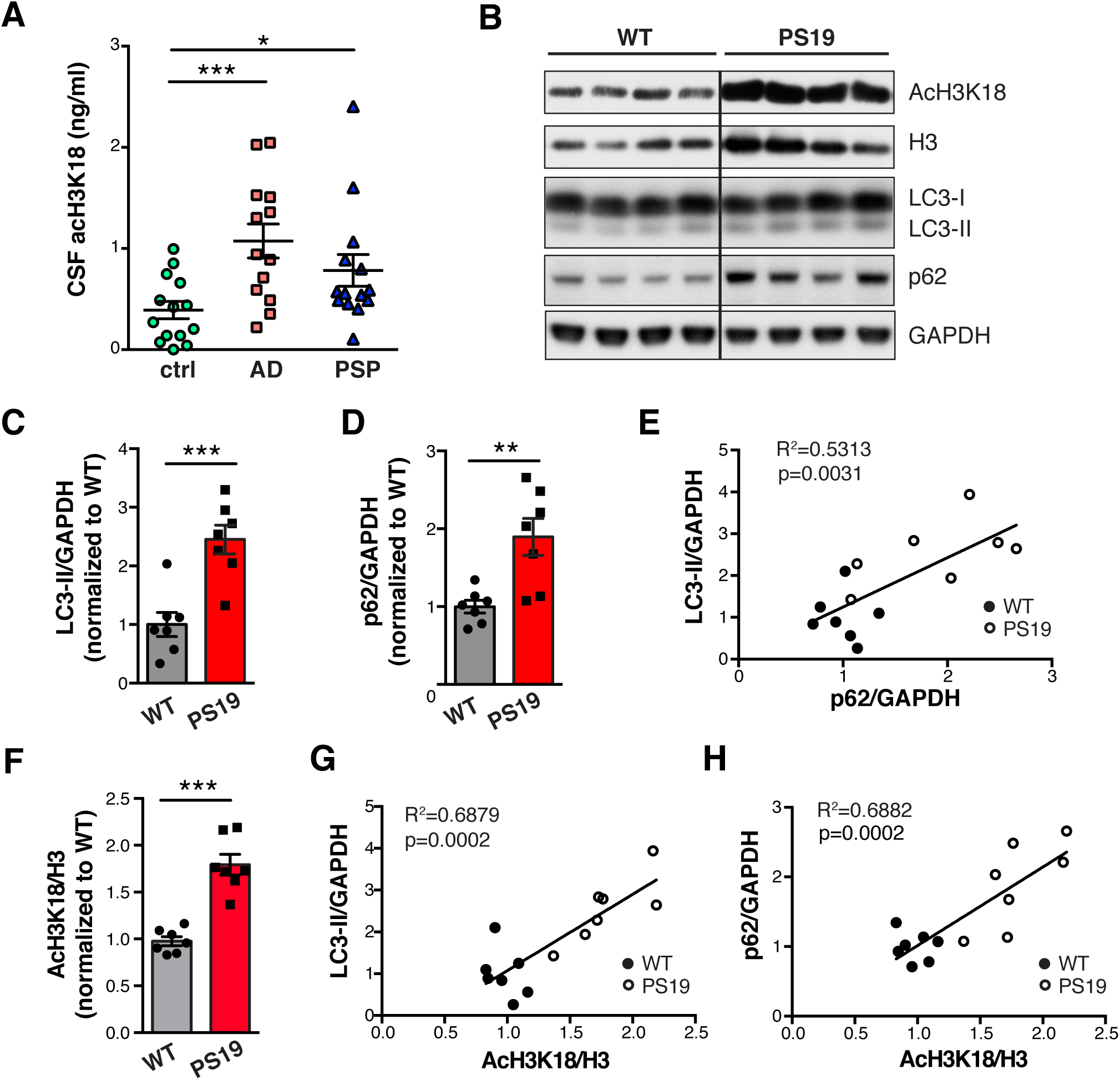
Hyperactive p300/CBP Is Associated with Impaired Autophagy in Tauopathy Brains. (A)Levels of acH3K18, measured by ELISA, in CSF samples from normal controls (ctrl, n = 14) and patients with AD (n=13) and PSP (n=14). *p<0.05, ***p<0.001 one-way ANOVA and Tukey-Kramer post hoc analysis. Values are mean ± (B)Representative immunoblots of acH3K18, total H3, LC3-I, LC3-II, SQSMT/p62, and GAPDH in hippocampal lysate of 10-month-old wildtype (WT) and PS19 mice. (C, D) Levels of LC3-II (C) and p62 (D) relative to GAPDH, normalized to WT. (E) Pearson correlation analysis of normalized LC3-II and p62 levels. AcH3K18 levels relative to total H3, normalized to WT. (G, H) Pearson correlation analysis of acH3K18 and LC3-II levels (G) and acH3K18 and p62 levels (H). (C–H) n=7 mice per group. **p<0.01, ***p<0.001 by unpaired *t* test. Values are mean ± SEM.

In AD brains with high burden of tau, autophagy is impaired, as shown by the accumulation of premature autophagic vacuoles (AVs), which are rare in healthy brains (Nixon et al., 2005, Boland et al., 2008). Because p300 has been reported to regulate autophagy (Lee and Finkel, 2009, Su et al., 2017), we investigated the relation between increased p300/CBP and impaired autophagy in tauopathy brains. We examined the autophagy markers LC3-II and SQSTM1/p62 in the hippocampi of 10-month-old PS19 mice, which had increased acH3K18 levels, indicating p300/CBP hyperactivity (Figures 1B and 1F). Levels of LC3-II and p62 were significantly higher in PS19 mice than in nontransgenic littermates (Figures 1B–1D). Since LC3-II and p62 are degraded in autolysosomes upon autophagosome maturation, higher steady-state levels of these markers suggests an accumulation of immature AVs in the brains, consistent with a previous immuno-electron microscopy study (Nixon et al., 2005). Indeed, levels of LC3-II and p62 correlated with each other (Figure 1E) and with p300/CBP activity (Figures 1G and 1H).

To pinpoint where active p300/CBP is localized along the ALP, we examined the localization of p300 and autophagic markers in tauopathy brains by immunohistochemistry. p-p300 (S1834), an active form of p300 (Huang and Chen, 2005, Wang et al., 2013), is abundant in cytosolic structures resembling granulovacuolar degeneration bodies (GVBs) in human AD brains (Aubry et al., 2015). GVB lesions are strongly associated with fibrillar tau pathology and are labeled by casein kinase 1 (CKiδ) (Ghoshal et al., 1999). In PS19 brains, neurons with AT8-positive tau inclusions were enriched in p-p300, which co-localized with CKiδ-positive GVB-like structures that overlapped with the late-endosomal marker chmp2B (Figures S2A–S2C). In contrast, nontransgenic mice were mostly negative for p-p300 and CKiδ puncta (Figures S2A′–S2C′). Interestingly, p-p300-positive granules were encircled by the endolysosomal marker lamp1 (Figure S2D). As p-p300 accumulates at late-stage autophagic intermediates in tau-accumulating neurons, we hypothesize that hyperactive p300/CBP impairs the ALP in tauopathy brains.

### p300/CBP Inhibits Autophagic Flux

To directly examine how p300/CBP hyperactivation affects autophagy in tau-bearing cells, we measured LC3-II and p62 levels in HEK293T cells overexpressing p300 or tau. Overexpression of p300 increased steady-state levels of both LC3-II and p62 (Figures 2A and 2B). Blocking lysosomal degradation with bafilomycin A1 (BafA1) led to accumulation of LC3-II and p62, as expected (Figure 2C), which allows for measurement of autophagic flux (Klionsky et al., 2012, Mizushima et al., 2010). In p300-overexpressing cells, the difference in LC3-II or p62 levels between DMSO and BafA1 treatment is significantly smaller, suggesting reduced autophagic flux (Figures 2C and 2D). In addition, a dual-labeled fluorescent reporter, mCherry-GFP-LC3, was used to monitor the maturation of AVs (Pankiv et al., 2007): immature AVs with neutral pH are yellow since both mCherry and GFP are fluorescent, whereas mature AVs with acidic pH are red since only mCherry retains fluorescence. In HEK293T cells expressing mCherry-GFP-LC3 alone or together with tau, more red AVs were observed than yellow AVs (Figures 2E and 2F), indicating that most AVs were mature and autophagic flux was normal. In contrast, overexpression of p300 significantly reduced the number of red AVs (Figures 2E and 2F) and the ratio of red to yellow AVs (Figure 2G), as well as the total number of AVs (Figure 2F). Thus, in cells expressing tau, increased p300 impairs autophagic flux both by reducing autophagosome induction and compromising maturation of autophagosomes into autolysosomes.

**Figure 2.**
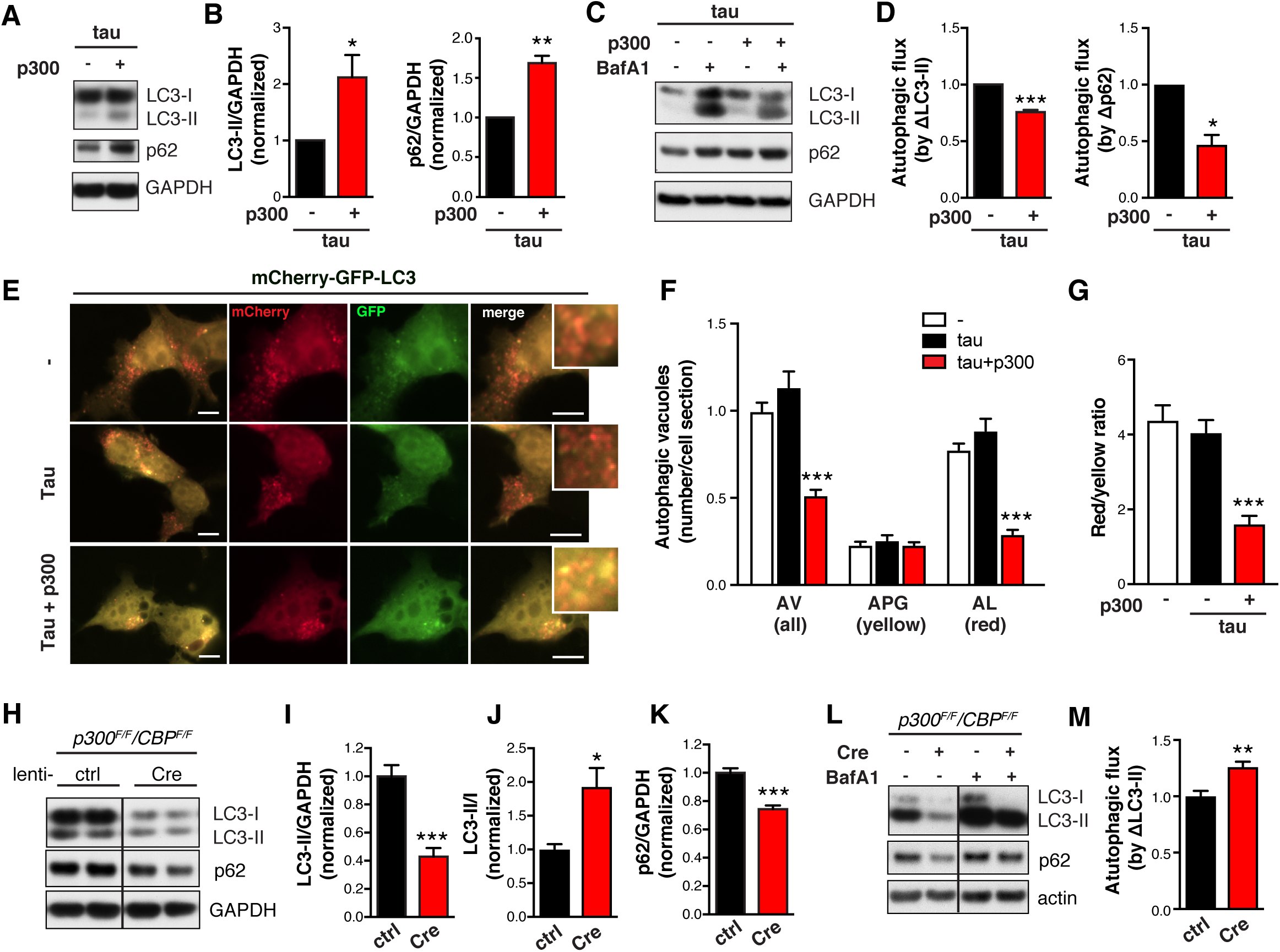
p300 Impairs Autophagic Flux. (A, B) p300 overexpression in HEK293T cells increases LC3 and p62 accumulation. (A) Representative immunoblot of LC3-I, -II, SQSMT/p62 and GAPDH in lysates of HEK293T cells transfected with tau alone or tau and p300 and serum-starved for 48 h. (B) Quantification of levels of LC3-II and p62 relative to GAPDH, normalized to control (tau alone). n=3 wells from three independent experiments. *p<0.05, **p<0.01 by unpaired *t* test. Values are mean ± SEM.(C, D) p300 overexpression in HEK293T cells reduces autophagic flux. (C) Representative immunoblots of LC3-I, -II, p62, and GAPDH in lysates of HEK293T cells transfected with tau alone or tau and p300 and treated with BafA1 (10 nM, 24 h) or DMSO in serum-free medium. (D) Quantification of autophagic flux by the change (increase) of LC3-II or p62 in response to BafA1, normalized to control (p300-, BafA1-). n=4 wells from two independent experiments. *p<0.05, **p<0.01, ***p<0.001 by unpaired *t* test. Values are mean ± SEM. (E–G) HEK293T cells stably expressing mCherry-GFP-LC3 are transfected with vector (-), tau, or tau+p300, and serum starved for 4 h to induce autophagy. (E) Representative images of mCherry and GFP fluorescent signal. Scale bar: 10 μm. (F) Quantification of autophagic vesicles (AV). Autophagosomes (APG) are identified as yellow vesicles retaining both mCherry and GFP fluorescence. Autolysosomes (AL) are identified as red vesicles in which GFP fluorescence is quenched by the low pH in lysosomes. ***p<0.001, two-way ANOVA, Tukey-Kramer post hoc analysis. (G) Ratio of the number of red vesicles to yellow vesicles per cell. ***p<0.001, one-way ANOVA, Tukey-Kramer post hoc analysis. (F, G) From two independent experiments, n=51 cells (-), n=24 cells (tau), and n=53 cells (tau+p300). Values are mean ± SEM. (H–K) p300/CBP double knockout in primary mouse neurons reduces LC3 and p62 accumulation. (H) Representative immunoblots of LC3-I, -II, p62, and GAPDH in lysates of *p300*^*F/F*^*/CBP*^*F/F*^ primary neurons infected with lentivirus expressing empty vector (lenti-ctrl) or Cre recombinase (lenti-Cre). (I–K) Quantification of LC3-II and p62 relative to GAPDH, and LC3-II/I, normalized to control. n=6 wells from three independent experiments. (L, M) p300/CBP double knockout in primary mouse neurons increases autophagic flux. (L) Representative immunoblots of LC3-I, -II, p62, and actin in lysates of p300/CBP double-knockout neurons that were treated with BafA1 (10 nM, 24 h) or DMSO. (M) Quantification of autophagic flux based on the change (increase) of LC3-II in response to BafA1 treatment, normalized to control (Cre-, BafA1-). n=4 wells from two independent experiments. *p<0.05, **p<0.01, ***p<0.001 by unpaired *t* test. Values are mean ± SEM.

We next examined the effects of p300/CBP inhibition on autophagic flux in neurons expressing human tau (hTau). Postmitotic knockout of p300 and CBP was done by infecting primary mouse neurons carrying loxP-flanked *Ep300* (*p300*^*F/F*^) and *CBP* (*CBP*^*F/F*^) with lentivirus expressing Cre recombinase. A lentiviral vector was used to transduce hTau into these neurons. Depletion of p300 and CBP was confirmed by the absence of p300, CBP, and acH3K18, as shown by immunoblot (Figure S1A, S1C). The double knockout did not affect neuronal viability or cytotoxicity (Figure S1B) but significantly reduced the levels of LC3-I, LC3-II, and p62 (Figures 2H, 2I, and 2K). Upon BafA1 treatment, p300/CBP knockout neurons and wildtype neurons accumulated similar amounts of LC3-II and p62 (Figure 2L). Thus, the reduced level of LC3-II in p300/CBP knockout neurons was due to enhanced autophagic flux rather than to reduced induction of autophagy (Figure 2M). Indeed, the conversion of LC3-I to LC3-II is accelerated (Figure 2J), indicating effective autophagic induction and flux in p300/CBP knockout neurons.

### p300/CBP Promotes Tau Secretion

Because it lacks a signal peptide, tau is thought to be released through unconventional secretion (Chai et al., 2012, Karch et al., 2012), which overlaps with the ALP (Mohamed et al., 2014, Zhang and Schekman, 2013, Ponpuak et al., 2015). Since p300 reduces autophagic flux, we investigated whether p300 hyperactivation affects tau secretion. In HEK293T cells overexpressing hTau, extracellular tau was readily detectable in the conditioned medium and was largely unphosphorylated (Figure 3A), consistent with previous reports (Pooler et al., 2013, Plouffe et al., 2012). As expected, co-expression of p300 markedly increased the level of acetylated tau (ac-tau), a considerable fraction of which was in the extracellular medium (Figure 3A). p300 overexpression increased intracellular accumulation of total tau (t-tau) (Figures 3A and S3A), as expected, since hyperacetylation of tau increases its stability and promotes its accumulation (Min et al., 2015, Min et al., 2010). Importantly, p300 overexpression led to significantly higher levels of extracellular tau relative to the intracellular tau (Figures 3B, S3A, and S3B), and markedly increased the levels of AT8-positive secreted phosphorylated tau (p-tau) without affecting levels of intracellular p-tau (Figures 3A, 3C, S3C, and S3D).

**Figure 3.**
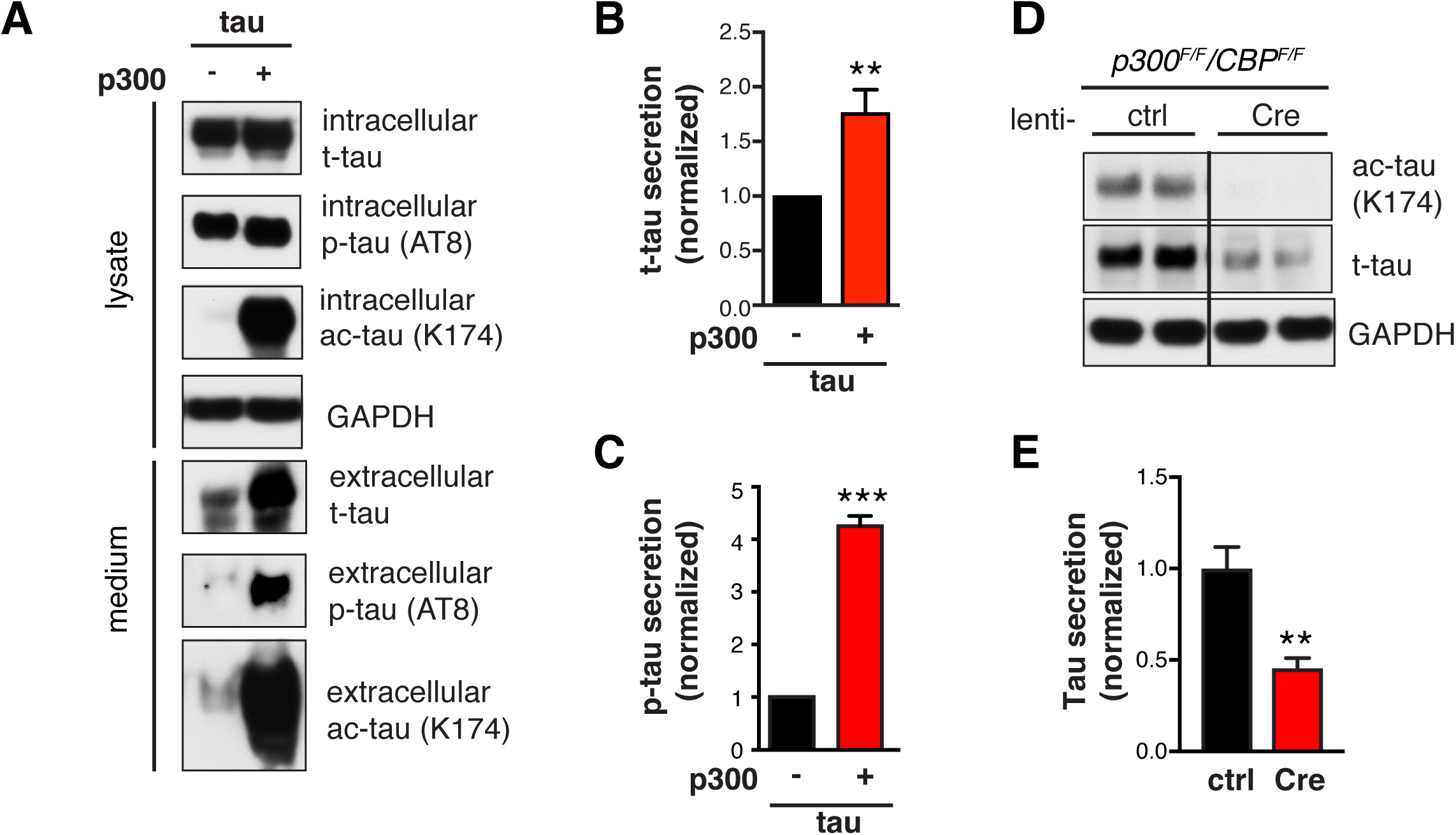
p300 Promotes Tau Secretion. (A–C) p300 overexpression in HEK293T cells increases tau secretion. (A) Representative immunoblot of total tau (t-tau), phosphor-tau (p-tau, AT8), acetylated tau (ac-tau, K174), and GAPDH in the lysate (5% of total) and conditioned medium (25% of total) of HEK293T cells transfected with tau alone or tau+p300. HEK293T cells were serum-starved for 48 h after transfection. The conditioned medium was concentrated 25-fold. (B, C) Quantification of t-tau secretion (B) and p-tau secretion (C) relative to intracellular levels, normalized to control. n=8 wells from eight independent experiments. **p<0.01, ***p<0.001 by unpaired *t* test. Values are mean ± SEM. (D, E) p300/CBP double knockout in primary mouse neurons reduces tau accumulation and tau secretion. (D) Representative immunoblots of ac-tau (K174), t-tau, and GAPDH in lysates of *p300*^*F/F*^*/CBP*^*F/F*^ primary neurons infected with lenti-ctrl or lenti-Cre. (E) Quantification of relative tau secretion over 3 h, based on levels of extracellular tau and intracellular tau measured by ELISA and normalized to control. n=6 wells from three independent experiments. **p<0.01, unpaired *t* test. Values are mean ± SEM.

To determine the role of endogenous p300/CBP in tau secretion, we generated p300/CBP double-knockout primary neurons, in which tau acetylation was abolished as expected (Figure 3D). p300/CBP ablation reduced intracellular t-tau levels by 50% without affecting tau mRNA, suggesting effects on protein turnover (Figures 3D, S3E, and S3F). Strikingly, p300/CBP knockout neurons secreted 75% less tau over 3 h than wildtype neurons (Figure S3G). Although part of the reduced secretion could be a consequence of accelerate intracellular turnover, the ratio of secreted to intracellular tau was significantly reduced in the knockout neurons (Figure 3E). Thus, p300/CBP, which inhibits autophagy flux, promotes tau secretion.

### A New p300 Inhibitor Reduces Tau Secretion

Next, we screened 133,000 compounds from the SMDC library for small-molecule inhibitors of p300, using a high-throughput homogeneous time-resolved fluorescence assay to measure the acetylation of full-length 2N4R tau by the purified core domain of p300 in the presence of acetyl-CoA (Figure 4A). Of these compound, 1120 had inhibition values >3 SD above the mean and were not promiscuous pan-assay interference compounds or quenchers of the donor. After confirming the hits and counter-screening to filter out fluorescence interference compounds, we confirmed 212 hits, of which 28 had an IC_50_<_50_ µM and a hillslope <3. Twenty of the 28 compounds were commercially available and purchased for retesting (Figure 4B). One of the top hits, 37892 (Figure 4C), had an IC_50_ of 35.3 μM (Figure S4A). Orthogonal MMBC assay confirmed its inhibitory activity (Figure S4B). 37892 competed with tau, as its IC_50_ values increased with increasing tau concentrations (Figure S4C). The binding of 37892 and p300 was determined by differential scanning fluorimetry (Figure S4D). In HEK239T cells, 37892 reduced p300 activity (measured by acH3K18) in dose-dependent fashion (Figures 4D and 4E) and did not induce significant toxicity (data not shown).

**Figure 4.**
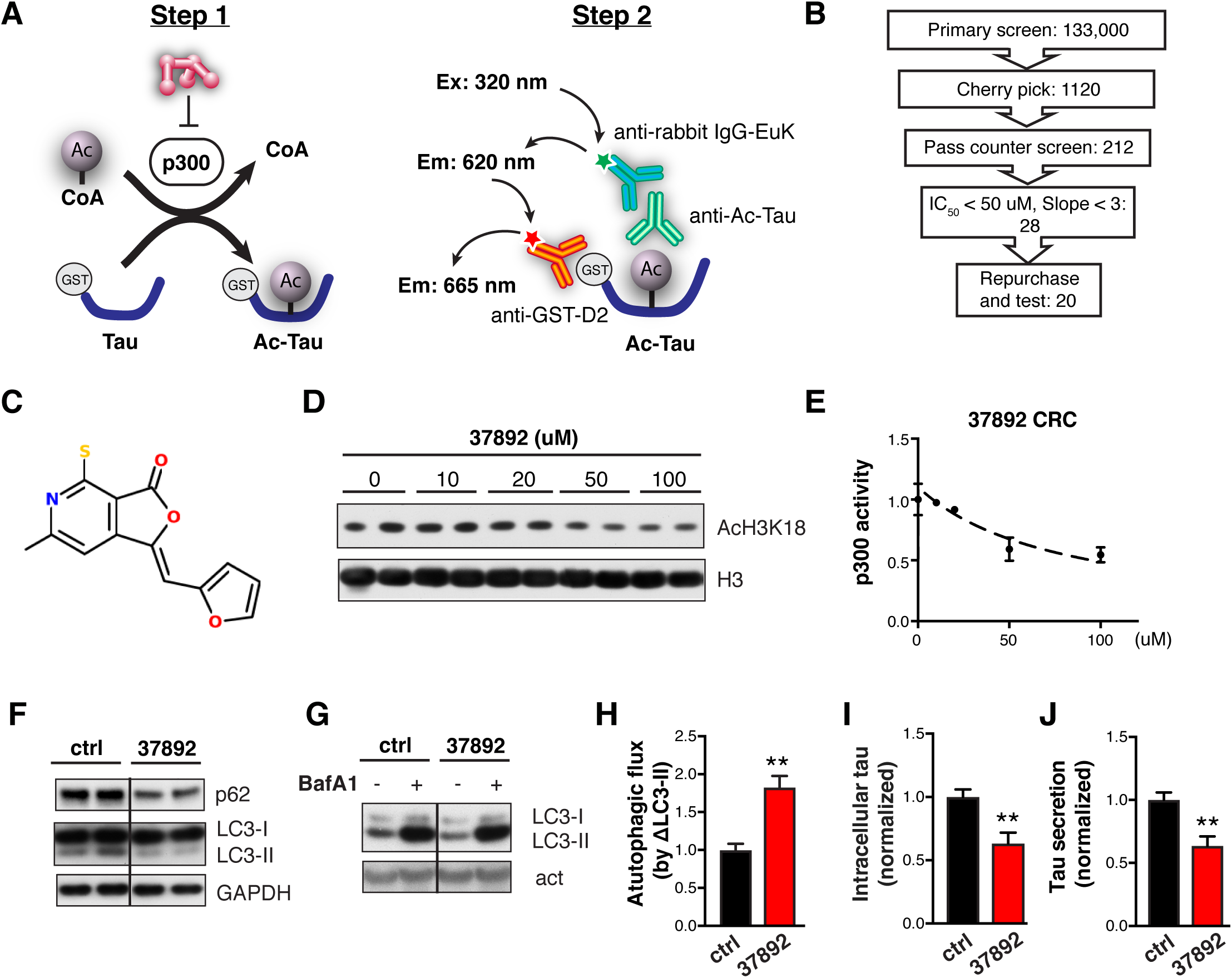
New p300 Inhibitor 37892 Reduces Tau Secretion. (A)Diagram of the homogeneous time-resolved fluorescence assay. Step 1, the enzymatic reaction: p300 transfers an acetyl group from acetyl-CoA to tau, producing CoA and ac-tau. Step 2, detection: a rabbit ac-tau–specific antibody (mAB359) recognizes ac-tau, the europium cryptate-labeled anti-rabbit IgG-EuK (donor) binds to mAB359, and the anti-GST-D2 (acceptor) captures GST-tau. When tau is acetylated, the Eu donor and D2 acceptor are brought into proximity, allowing Forster resonance energy transfer (FRET) to occur. FRET is detected as a long-lived fluorescence signal at 665 nm. The fluorescence signal ratio 665/620 nm is proportional to the extent of ac-tau in the solution. (B)Work flow of the high-throughput screen. (C)Structure of SMDC37892. (D, E) 37892 inhibits p300 activity in HEK293T. (D) Representative immunoblots of acH3K18 and H3 in histone extracts of HEK293T cells treated with increasing doses of 37892 for 24 h. (E) Concentration-response curve for 37892, quantified from p300 activity in HEK293T cells. (F–J) 37892 treatment in hTau-expressing rat primary neurons. (F) Representative immunoblots of p62, LC3-I, -II and GAPDH in primary neurons treated with 37892 (50 μM) for 24 h. (G) Representative immunoblots of LC3-I, -II and actin in primary neurons treated with 37892 (50 μM), BafA1 (5 nM), or both, for 24 h. (H) Autophagic flux in control and 37892-treated neurons quantified from the increase in LC3-II levels in response to BafA1 treatment, normalized to control. (I) Quantification of intracellular tau levels by ELISA and normalized to control. (J) Quantification of relative tau secretion over 3 h, based on levels of extracellular tau and intracellular tau measured by ELISA and normalized to control. n=6 wells from three independent experiments. **p<0.01 by unpaired *t* test. Values are mean ± SEM.

We next examined the effects of 37892 on autophagy and tau homeostasis in primary neurons overexpressing hTau. 37892 reduced LC3-II and p62 accumulation under steady state (Figure 4F), and increased autophagic flux as measured by changes in LC3-II level after co-treatment with BafA1 (Figures 4G and 4H). On the other hand, 37892 significantly reduced tau accumulation and tau secretion (Figures 4I and 4J). Similarly, in human iPSC-derived excitatory neurons (Wang et al., 2017a), 37892 reduced p300 activity (Figures S5B, S5F and S5G), reduced intracellular ac-tau and t-tau accumulation (Figures S5A, S5C, S5D and S5F), and reduced tau secretion (Figure S5E). To determine if 37892’s effect on tau secretion is dependent on p300 inhibition, we treated p300/CBP double knockout neurons with 37892, and failed to see further decrease of tau secretion in addition to the effect of knockout alone (Figure S5H). Therefore, 37892 inhibits tau secretion through inhibition of p300.

### Autophagic Flux Modulates Tau Secretion

To investigate whether changes in autophagic flux affect tau secretion, we blocked lysosomal degradation with a combination of NH_4_Cl (pH neutralizer) and leupeptin (inhibitor of lysosomal enzymes) (N/L). In hTau-expressing primary neuron, N/L treatment led to significant accumulation of LC3-II and p62, as expected (Figures 5A–5C). The intracellular tau level was increased (Figure 5D), indicating that tau is normally degraded by the ALP. Remarkably, relative tau secretion was increased by two-fold (Figure 5E). The increase was not due to cell death or increased membrane permeability, as lactate dehydrogenase (LDH) release was unaffected (Figure S6A). Similar effects were observed in primary neurons treated with BafA1 (Figures S6B–S6F). Treatment with N/L (Figures S6G, S6H), or vinblastine—which inhibits the fusion of autophagosomes with endosomes and lysosomes (Kochl et al., 2006)—also promoted secretion of endogenous tau in human iPSC-derived neurons (Figures S6I, S6J). In complementary experiments to investigate whether increasing autophagic flux reduces tau secretion, treatment with rapamycin—which induces autophagy by inhibiting mammalian target of rapamycin (Jung et al., 2010)—increased LC3-II and reduced p62 in primary neurons (Figures 5F–5H) and significantly reduced tau secretion (Figure 5J). Thus, tau secretion is strongly influenced by autophagic flux.

**Figure 5.**
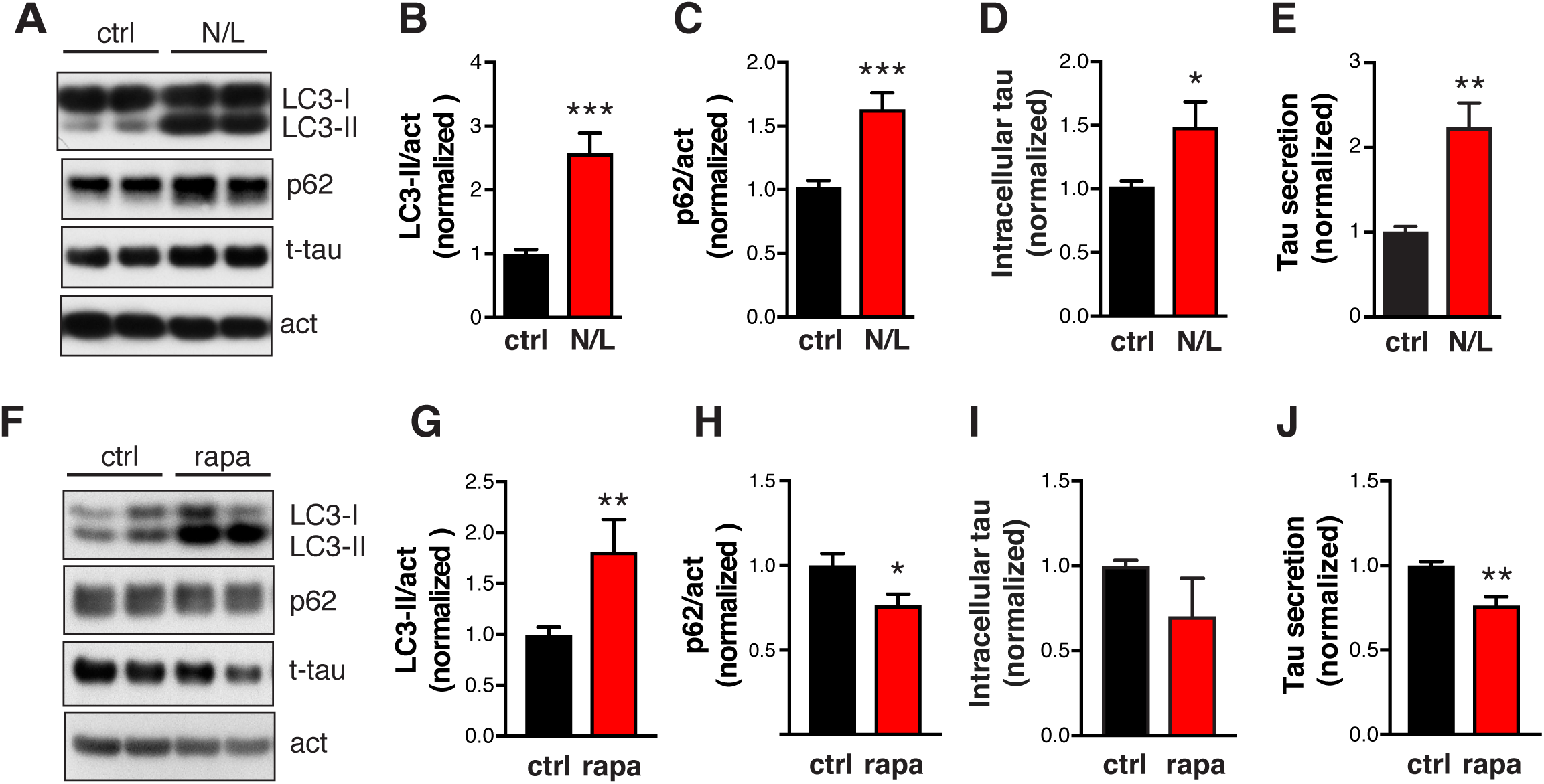
Autophagic Flux Modulates Tau Secretion. (A–E) Blocking autophagic flux with NH^4^Cl/leupeptin (N/L) increases tau secretion in primary neurons expressing hTau. (A) Representative immunoblots of LC3-I, LC3-II, p62, and actin in lysates of rat primary neurons infected with AAV-P301S hTau, after 24 h treatment with ctrl (DMSO) or N/L (20 mM NH^4^Cl and 200 μM leupeptin). (B, C) Quantification of levels of LC3-II and p62 relative to actin, normalized to control. (D) Quantification of intracellular tau levels by ELISA and normalized to control. (E) Quantification of relative tau secretion over 3 h, based on levels of extracellular tau and intracellular tau measured by ELISA and normalized to control. n=6 wells from three independent experiments. ***p<0.001, **p<0.01, *p<0.05 by unpaired *t* -test. Values are mean ± SEM. (F–J) Enhancing autophagic flux with rapamycin (rapa) reduces tau secretion in primary neurons expressing hTau. (F) Representative immunoblots of LC3-I, LC3-II, p62, and actin in lysates of rat primary neurons infected with AAV-P301S hTau, after 24 h of treatment with DMSO (control) or rapa (0.5 μM). (G, H) Levels of LC3-II (G) and p62 (H) relative to actin, normalized to control. (I) Quantification of intracellular tau levels by ELISA and normalized to control. (J) Quantification of relative tau secretion over 3 h, based on levels of extracellular tau and intracellular tau measured by ELISA and normalized to control. n=4 wells from two independent experiments. **p<0.01, *p<0.05 by unpaired *t* test. Values are mean ± SEM.

### p300/CBP Promotes Tau Secretion by Reducing Autophagic Flux

Since p300/CBP inhibits autophagic flux and promotes tau secretion, we next asked whether p300/CBP-induced excessive secretion of tau and p-tau is due to impairment of autophagic flux. We tested how autophagy affect tau secretion induced by p300 overexpression, and vice versa. In the absence of p300 overexpression, rapamycin enhanced autophagic flux, as indicated by an increased turnover of LC3-I to LC3-II (reflected in reduced LC3-I levels) and reduced p62 levels (Figures 6A and S6K), and reduced t-tau secretion (Figures 6A and 6B). Enhancing autophagy flux by rapamycin significantly reduced the secretion of t-tau, p-tau, and ac-tau induced by p300 overexpression (Figures 6A–6C). We then examined whether the reduction in tau secretion associated with p300 inhibition is mediated through autophagy. Primary neurons expressing hTau were co-treated with p300 inhibitor 37892 and lysosomal inhibitor BafA1. 37892 treatment alone reduced tau secretion, as expected (Figure 6D). When neurons are co-treated with 37892 and BafA1, tau secretion is elevated to similar levels as BafA1 treatment alone (Figure 6D). Thus, blocking autophagic flux by BafA1 abolished 37892’s effects on lowering tau secretion. Finally, we examined whether the effect of p300 hyperactivity and ALP inhibition are additive. While p300 overexpression alone and BafA1 treatment alone increased tau secretion in HEK293T, the combination of the two did not further increase tau secretion (Figure 6E). Together, these results suggest that the effect of ALP impairment on tau secretion is downstream of p300/CBP.

**Figure 6.**
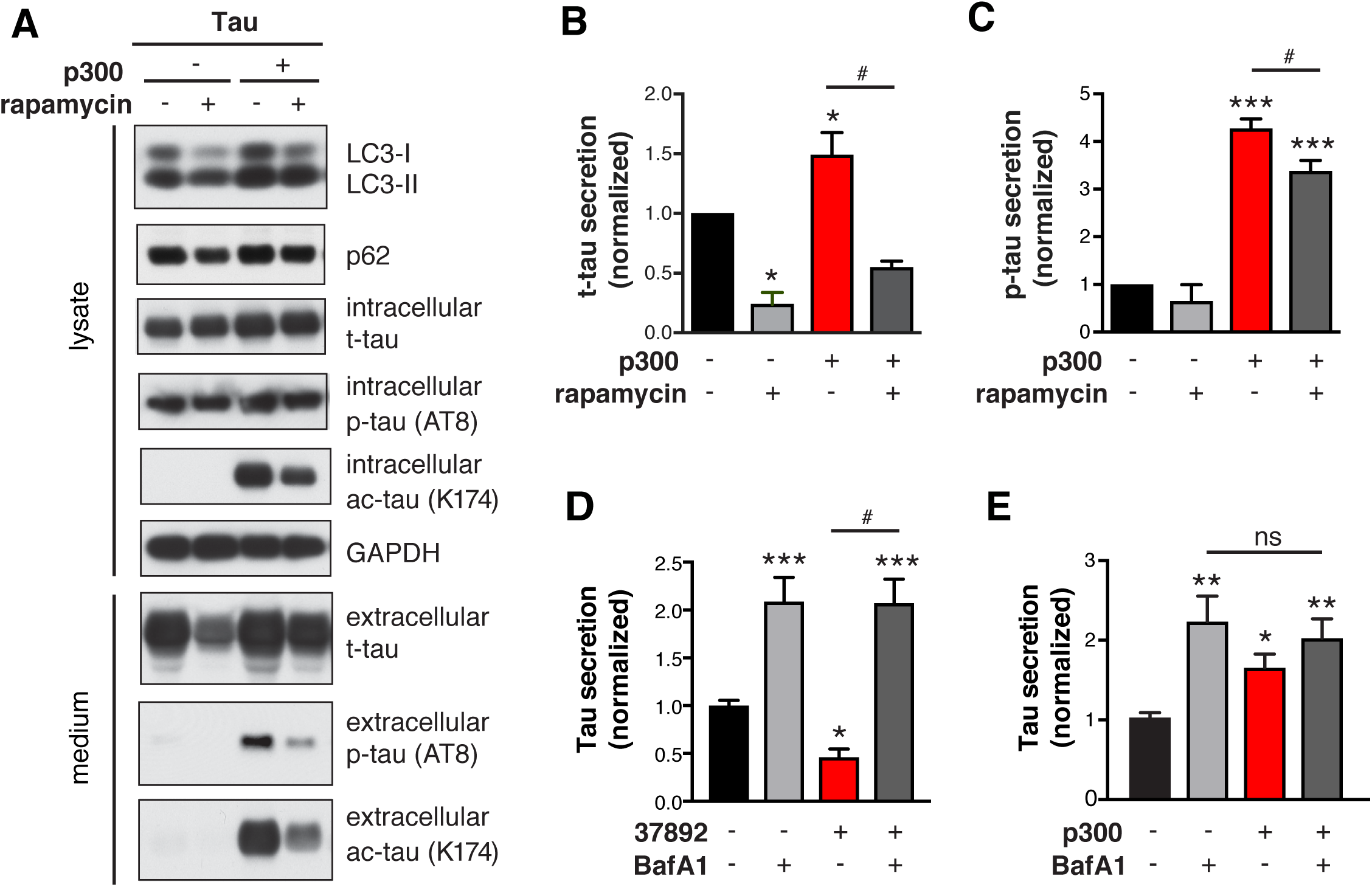
p300 Promotes Tau Secretion by Reducing Autophagic Flux. (A–D) Rapamycin treatment in HEK293T cells with and without p300 overexpression. (A) Representative immunoblots of LC3-I, LC3-II, p62, total tau (t-tau), phosphor-tau (p-tau, AT8), acetylated tau (ac-tau, K174), and GAPDH in lysate (5% of total) and conditioned medium (25% of total) of HEK293T cells transfected with tau alone or tau + p300. Serum-starved HEK293T cells were treated with DMSO (ctrl) or rapamycin (1 μM) for 24 h. The conditioned medium was concentrated 25-fold. (B, C, D) Quantification of t-tau secretion (B) and p-tau secretion (C) relative to intracellular levels and normalized to control. n=3 wells from three independent experiments. *^, #^p<0.05, ***p<0.001 by one-way ANOVA and Sidak’s multiple comparisons test. Values are mean ± SEM. (D)Blocking autophagic flux with BafA1 abolishes the effect of 37892 on lowering tau secretion in primary neurons. Quantification of relative tau secretion in 3 h window in neurons treated with BafA1, 37892 or combined, based on levels of extracellular tau and intracellular tau measured by ELISA and normalized to control. n=6 wells from three independent experiments. *p<0.05, ***p<0.001 by one-way ANOVA and Sidak’s multiple comparisons test. Values are mean ± SEM. (C)Blocking autophagic flux with BafA1 occluded the effect of p300 overexpression on tau secretion in HEK293T cells. Quantification of relative tau secretion in 24 h relative to intracellular levels and normalized to control. n=7 wells from three independent experiments. *p<0.05, **p<0.01 by one-way ANOVA and Sidak’s multiple comparisons test. Values are mean ± SEM.

### Inhibition of p300/CBP Reduces Tau Spreading

Tau secretion from neurons in which pathogenic tau is accumulating is the first step toward intercellular transmission of tau and the spread of tau pathology. To determine how p300/CBP affects the spread of pathogenic tau, we used an in vitro model in which synthetic tau fibrils (K18/PL) induce the formation and intercellular spreading of tau aggregates in primary neurons expressing P301S hTau (Guo and Lee, 2013). MC1 immunostaining is used to detect the pathological, aggregated conformation of tau (Jicha et al., 1997) (Figure 7A). In *p300*^*F/F*^*/CBP*^*F/F*^ primary neurons, p300/CBP double knockout with lenti-Cre abolished the acH3K18 signal without affecting neuron number (Figures S1C and 7C). p300/CBP knockout markedly reduced the number of neurons bearing MC1-positive tau aggregates (Figures 7A and 7B). We also examined effect of p300/CBP heterozygous double knockout in *p300*^*F/+*^*CBP*^*F/+*^ neurons infected with lenti-Cre, in which partial inhibition of p300 and CBP significantly reduced the MC1 signal without changing neuron number and neurite length (Figures S7A-S7E). A similar effect was observed in primary neurons treated with 37892 (Figures S7F–7I). Thus, inhibition of p300/CBP reduces tau spreading in vitro.

**Figure 7.**
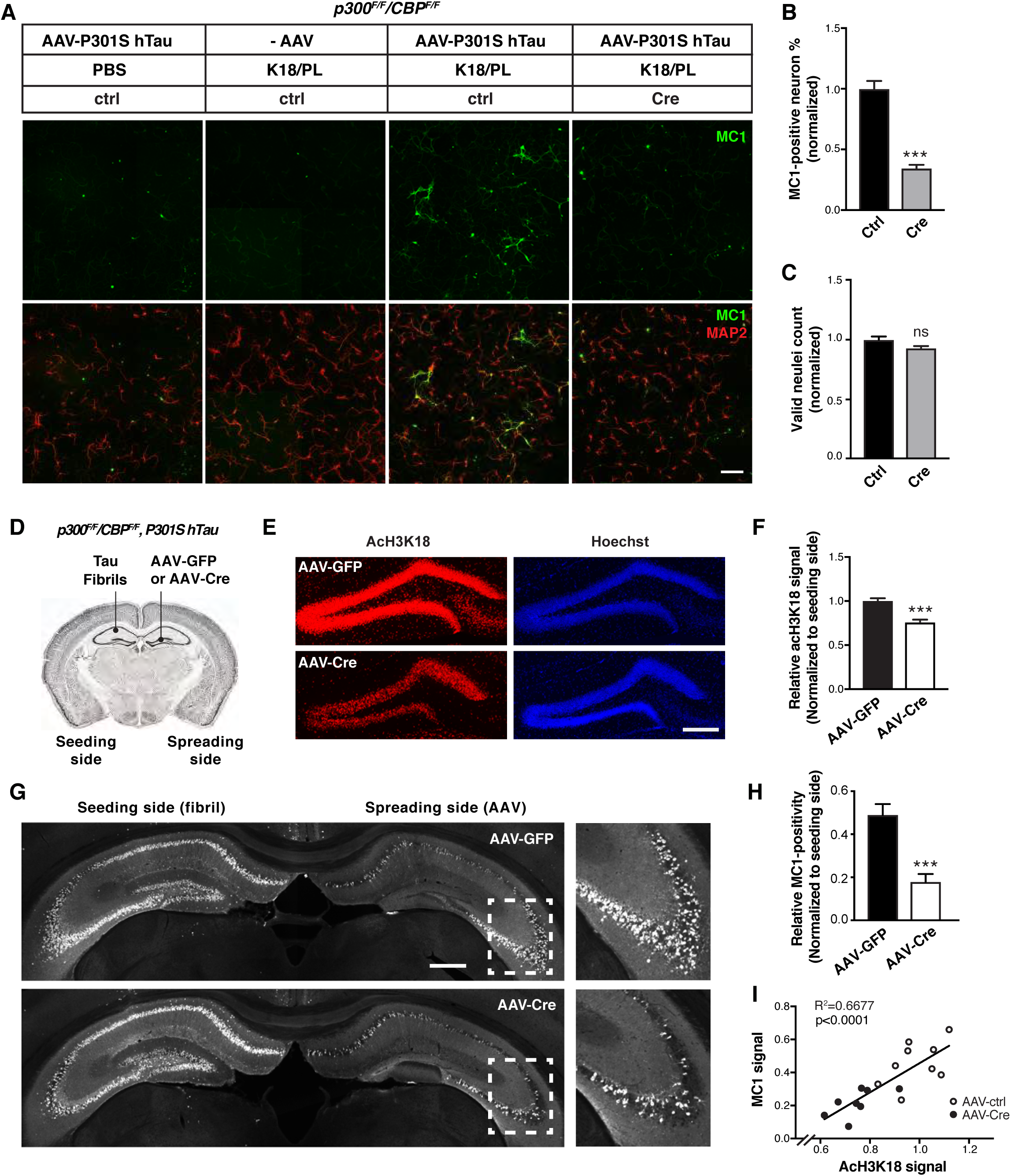
Inhibition of p300/CBP Reduces Tau Spreading. (A–C) p300/CBP double knockout reduces fibril-induced tau spreading in primary neurons. (A) Representative immunofluorescence staining with MC1 and MAP2 antibody in *p300*^*F/F*^*/CBP*^*F/F*^ primary neurons infected with AAV-P301S hTau and lenti-control or lenti-Cre, and treated with synthetic tau fibril (K18/PL, 100 nM). Negative controls (PBS-treated, AAV non-infected) are included. Scale bar: 100 μm. (B) Percentage of MC1-positive neurons, normalized to control. ***p<0.001, unpaired *t* test. (C) Number of valid (live) neuclei, normalized to control. n=9 wells from two independent experiments. Values are mean ± SEM. (D–I) Inhibition of p300/CBP reduces tau spreading in PS19 mice. (D) Schematic diagram of stereotaxic injections in the hippocampus of PS19 mice. Tau fibrils (K18/PL) were injected into left CA1 (seeding side). AAV-Cre or AAV-GFP were injected into right dentate gyrus (spreading side). (E) Representative images of immunostaining with acH3K18 antibody in the hippocampus after AAV-GFP (control) and AAV-Cre injections. Scale bar: 200 μm. (F) Quantification of acH3K18-positive area (normalized to Hoechst) on the spreading side (AAV-injected) normalized to the fibril-injected side of hippocampus. (G) Representative images of immunostaining of MC1 in the hippocampus after AAV-GFP and AAV-Cre injections. Scale bar: 500 μm. (H) Quantification of MC1-positive area on the spreading side (AAV-injected) normalized to the fibril-injected side of hippocampus. (I) Pearson correlation analysis of MC1 signal and acH3K18 signal (normalized). n = 7 slices from 9 (AAV-GFP) or 8 (AAV-Cre) mice per group. (F, H) ***p < 0.001, unpaired *t* test. Values are means ± SEM.

To determine whether p300/CBP inhibition reduces tau spreading in vivo, we simultaneously injected tau fibrils into the left hippocampus and AAV-Cre or AAV-GFP into the right hippocampus of 4–5-month-old PS19 mice carrying *Ep300*^*F/F*^and *CBP*^*F/F*^ (Figure 7D). AAV-Cre inhibited p300/CBP in the right hippocampus, as shown by reduced staining for acH3K18 (Figures 7E and 7F). Consistent with previous studies (Iba et al., 2013), injection of tau fibrils markedly increased MC1-positive tau pathology in the ipsilateral and contralateral hippocampi within 1 month (Figure 7G). Notably, Cre-mediated p300/CBP inhibition significantly reduced the spread of MC1-positive tau to the contralateral hippocampus (Figures 7G and 7H). Remarkably, the extent of tau spreading quantified by MC1 positivity correlated with p300/CBP activity, measured by acH3K18 signal (Figure 7I). Thus, inhibition of p300/CBP reduced the spread of tau pathology in vivo.

## DISCUSSION

This study shows that p300/CBP acetyltransferase activity is increased in human AD brains and correlates with impairment of autophagy in tauopathy mouse brains. In cultured cells and human iPSC-derived neurons, inhibition of p300/CBP enhanced autophagic flux and suppressed tau secretion and vice versa. Moreover, p300/CBP inhibition strongly reduced the pathogenic spread of tau inclusions in vivo. Through detailed mechanistic dissections, we showed that p300/CBP promotes tau secretion by inhibiting autophagic flux. These findings establish a novel connection between p300/CBP-mediated autophagy inhibition and tau propagation.

p300/CBP, best known as transcriptional cofactor, is a master regulator of AD progression (Aubry et al., 2015). We found that p300 activity is increased in the CSF of patients with AD and PSP, consistent with previous studies (Wong et al., 2013, Aubry et al., 2015). We previously showed that p300/CBP is the lysine acetyltransferase for tau, whose hyperactivation leads to increased tau acetylation and tau accumulation (Min et al., 2015, Min et al., 2010). How p300/CBP becomes hyperactive in diseases brain is unclear. Interestingly, mouse brains expressing high levels of mutant hTau had elevated levels of p300/CBP activity, which could exacerbate tau accumulation, resulting in a feedforward vicious cycle.

The elevated p300/CBP activities correlated with the impairment in autophagy in tauopathy mouse models. The intracellular homeostasis of tau requires proper degradation by the UPS and ALP (reviewed in (Lee et al., 2013). Autophagy-mediated tau degradation could be inhibited by pathogenic point mutations of tau (P301L, A152T) and post-translational modifications (Caballero et al., 2017). In AD brains, autophagic and endolysosomal dysfunction was accompanied by accumulation of immature autophagic vacuoles (Sanchez-Varo et al., 2012, Nixon et al., 2005). The mechanisms of autophagy impairment in late-onset sporadic AD are unclear. Our findings implicate p300/CBP hyperactivation as a key player that impairs autophagic flux and slows the turnover of autophagic vacuoles through lysosomal degradation. As previously reported in Hela cells, p300 acetylates Atg5, Atg7, Atg8, and Atg12, (Lee and Finkel, 2009), which affect autophagy flux. Importantly, inhibition of p300/CBP in primary neurons increased autophagic flux under normal nutrient condition. In our p300/CBP knockout neurons, LC3-II and p62 levels were both decreased, while LC3-II/I ratio was increased, suggesting robust autophagic flux. By contrast, in HeLa cells, knockdown of p300 reduced p62 levels and increased LC3-II levels (Lee and Finkel, 2009). This discrepancy is likely due to the difference in cell type–specific basal levels of autophagy and to a more efficient clearance of newly forming autophagosomes by lysosomes in the case of neurons (Boland et al., 2008). Nonetheless, it is consistent that p300/CBP inhibition increases autophagic flux among different cell types.

Besides inhibiting ALP, p300/CBP activation promoted secretion of tau, especially p-tau, by regulating autophagy activity. Blocking lysosomal degradation or the fusion of autophagosomes with lysosomes markedly increased tau secretion, whereas promoting autophagic flux with rapamycin reduced tau secretion. Thus, tau secretion could be an alternative clearance mechanism to maintain overall cellular proteostasis when the degradative pathways are impaired or overloaded. Autophagy-mediated unconventional secretion has been characterized for IL1β and other proteins that are secreted without the involvement of Golgi-ER complex (Zhang and Schekman, 2013, Ponpuak et al., 2015). In Parkinson’s disease, α-synuclein is secreted through a similar mechanism, where accumulated autophagic intermediates such as autophagosome, autolysosome and multivesicular bodies (MVBs) fuse with the plasma membrane and release α-synuclein in exosomes (Ejlerskov et al., 2013). Importantly, our findings suggest that p300/CBP affects tau secretion through effects on ALP, since increasing autophagic flux partially rescued p300/CBP-induced increase in tau secretion and vice versa. It is of interest to identify the downstream mediator in ALP for p300/CBP-induced tau secretion. As recently reported, p300-mediated acetylation of beclin 1, VPS34, and SIK2 inhibits autophagy initiation, autophagosome maturation, and endocytic trafficking (Sun et al., 2015, Su et al., 2017, Yang et al., 2013), suggesting that these proteins have a potential role in regulating tau secretion. Finally, as p300/CBP regulates histone acetylation, the mechanism could also involve epigenetic changes in the autophagy-relevant transcriptome (Eisenberg et al., 2014).

The exact subcellular compartment through which tau is secreted is unknown. We found that increased tau secretion is associated with accumulation of LC3-II and p62, markers of intermediated AVs. However, as previously reported, LC3-labeled autophagosomes do not appear to be enriched in tau (Mohamed et al., 2014); Similarly, we found tau inclusions largely reside in the cytosol without specific colocalization with a certain type of autophagic vesicles. Interestingly, in Atg5 knockout neurons lacking LC3-II-positive autophagosomes, we found that tau secretion was not abolished (data not shown), indicating that autophagosomes are dispensable for tau secretion. Tau has been found in exosomes isolated from cultured neurons and CSF, suggesting MVB-mediated release of exosomes is involved in tau secretion (Wang et al., 2017b, Ngolab et al., 2017). It was also reported that Rab7A, a small GTPase involved in endosomal trafficking, regulates tau secretion, indicating that a late endosomal compartment is involved (Rodriguez et al., 2017). Finally, as the last step in the ALP, lysosomes can undergo exocytosis, a process resembling synaptic vesicle release regulated by TFEB (Medina et al., 2011), which could also mediate tau secretion when autophagy is impaired. In support of the hypothesis that later-stage intermediate vesicles in the ALP mediates tau secretion, we found accumulation of GVB-like structures with molecular signature of late endosome and lysosome in P301Stau transgenic mouse brain. Various pathogenic tau species differ in their effects on different autophagy pathways (macroautophagy, chaperone-mediated macroautophagy and endosomalmicroautophagy) (Caballero et al., 2017). Complex crosstalk between autophagy pathways and between autophagy and endolysosomal pathways (Kaushik et al., 2008) can provide compensatory mechanisms for the interplay between tau degradation and secretion. For example, autophagy induction promotes the fusion of MVBs with lysosomes and inhibits exosome release (Fader et al., 2008). In line with this, we found that rapamycin induces autophagy and reduces tau secretion, both in wildtype cells and in cells overexpressing p300. Rab GTPases (e.g., Rab1A, Rab8, Rab7, and Rab27A) regulate vesicular transport and fusion events along the biosynthetic and endosomal pathway and differentially regulate α-synuclein secretion (Ejlerskov et al., 2013). Thus, fine-tuning of the ALP at multiple steps can push the system to reach a new balance between degradation and secretion of the cargo. Increased secretion of tau into the extracellular space is the basis for tau propagation to different brain areas, the spread of pathology, and disease progression. p300/CBP inhibition significantly reduces fibril-induced spreading of tau pathology, in which tau seeds secreted from cells are taken up by recipient cells, where they induce tau pathology. Here we showed that p300/CBP inhibition reduced both endogenous tau accumulation and the spread of aggregated tau pathology. Promoting autophagy by rapamycin reduces tau pathology in mouse models (Majumder et al., 2011, Ozcelik et al., 2013) and reduces tau spreading in a neuronal culture model (Xu et al., 2016). Our results suggest that inhibition of p300/CBP promotes autophagic flux, enhances tau degradation and reduces tau secretion, thereby attenuates tau spreading. p300/CBP inhibition also reduces tau acetylation, which in turn enhances tau clearance and reduces tau accumulation. Thus, the attenuated spread of tau pathology we observed with inhibition of p300/CBP might result from reduction of both intracellular tau accumulation and extracellular tau secretion. Tau propagation is thought to be caused by soluble tau species, either monomeric or oligomeric (Michel et al., 2014, Mirbaha et al., 2015). However, it remains to be determined whether p300/CBP affects posttranslational modification of transmitted tau species, or tau seed uptake. Finally, glial cells may contribute to tau spreading (Asai et al., 2015), and the p300/CBP-autophagy pathway could also be involved.

The link between p300/CBP and autophagy regulation highlights the important crosstalk between metabolic status and proteolysis system. The substrate of p300/CBP, acetyl-coenzyme A (AcCoA), serves as a major integrator and messenger of nutritional status. Increased or reduced cytosolic AcCoA suppresses or induces autophagy, respectively, mediated through p300/CBP (Marino et al., 2014). On the other hand, the effects of acetylation on autophagy are reversed by SIRT1 or HDAC6-mediated deacetylation (Lee et al., 2008, Huang et al., 2015, Sun et al., 2015, Yang et al., 2013, Garcia-Aguilar et al., 2016, Lee et al., 2010), which is also regulated by the availability of nutrition for the cell.

In summary, our findings provide direct evidence to link p300/CBP hyperactivation, ALP impairment and tau secretion and propagation. This mechanistic link suggests that interfering with p300/CBP-autophagy pathway may be a promising therapeutic strategy to counteract tau pathogenesis.

## ACKNOWLEDGMENTS

We thank Drs. E. Verdin and E. Koo for insightful discussions, Dr. P. Davies (Feinstein Institute for Medical Research) for MC1 antibody, Drs. S. Ding and M. Xie for medicinal chemistry advice; D. Le, Y. Zhou and M. Chin for technical assistance, members of the Gan lab for discussions, E. Nguyen for administrative assistance, and Stephen Ordway for editorial support. This work was supported by a grant from Rainwater Foundation (to L.G.), US National Institutes of Health (NIH) grants U54NS100717 and R01AG054214 (to L.G.), R01NS059690 (to J.E.G.) and K99 AG053439 (to X.C.).

## AUTHOR CONTRIBUTIONS

L.G. and X.C. conceived the project. L.G., X.C. and M.A. designed experiments. X.C., Y.L., C.W., and Y.T. performed experiments. S-A.M., R.M.T., J.C.R., A.K., B.L.M., A.L.B., J.E.G. and A.M.C. developed experimental tools or reagents. X.C., L.G., Y.T. and A.M.C. wrote the manuscript.

## EXPERIMENTAL PROCEDURES

### Primary Antibodies and Chemicals

Acetylated H3/K18 (Abcam), H3 (Cell Signaling), LC3B (Acris), LC3A (Novus Biologicals), p62/SQSTM1 (Novus Biologicals), GAPDH (Sigma), actin (DSHB), Tau-5 (Bio-Source), HT7 (Thermo Fisher), BT2 (Thermo Fisher), AT8 (Thermo Fisher), MAP2 (Millipore), p-p300 (S1834) (Sigma), CKiδ (Abcam), Chmp2B (Abcam), and lamp1 (1D4B, DSHB), Europium cryptate-labeled anti-rabbit IgG-EuK (61PARKLB, Cisbio) and anti-GST-D2 (61GSTDLB, Cisbio), were purchased. Ac-tau/K174 antibody (AC312) and ac-tau/K274 antibody (mAB359) were produced and characterized as described previously(Min et al., 2015, Tracy et al., 2016). MC1 antibody is a gift from Dr. Peter Davies (Feinstein Institute). NH4Cl (Sigma), Leupeptin (Sigma), vinblastine (Sigma), rapamycin (Invivogen), and Bafilomycin A1 (Invivogen) were purchased.

### Human Samples

CSF samples were obtained from 13 patients with Alzheimer’s disease (AD), 18 patients with progressive supranuclear palsy (PSP) and 14 cognitively normal participants. Patients were recruited through the research programs: Frontotemporal Dementia: Genes, Images and Emotions, 4-Repeat Tauopathy Neuroimaging Initiative (4RTNI), Alzheimer’s Disease Research Center (ADRC) and a PSP clinical trial (NCT02422485) at the University of California, San Francisco (UCSF). All patients received a comprehensive neurological history, physical, structured caregiver interview and neuropsychology assessment. Diagnosis was made by consensus panel, utilizing the standard diagnostic criteria for AD and PSP (Litvan et al., 1996, McKhann et al., 2011). Cognitively normal healthy patients were recruited through the Larry L. Hillblom Aging Study at UCSF. The eligibility criteria included ages 60-100, with no significant subjective memory complaints, no functional impairment, a Clinical Dementia Rating (CDR) of 0, no diagnosis of mild cognitive impairment, and a Mini-Mental State Exam (MMSE) score of ≥ 26. All participants provided written and informed consent, and the institutional review board of UCSF approved all studies.

CSF collection and processing was performed according to the Alzheimer’s Disease Neuroimaging Initiative protocol. CSF was obtained by lumbar puncture using a 25-gauge needle and collected in 10-mL polypropylene tubes. Within 1 h, CSF was centrifuged at 2,000g for 10 minutes at 4°C, transferred to new polypropylene tubes and stored at −80°C until analysis.

### Protein Purification

DNA encoding a fragment of p300 (aa 1815-1910), including the PHD, catalytic (HAT), ZZ and TAZ2 domains, was cloned into a pET24b vector. The plasmid was transformed into the E. coli Rosetta BL21 strain (Invitrogen). Frozen cell stock was streaked onto Kanamycin (50 μg/mL) plate and grown overnight. One colony was picked and grown in a starter culture and used to inoculate 6 L of 2X YT media. Upon log-phase growth (OD ∽0.6-0.8), expression was carried out by overnight induction with 0.2 mM IPTG at 16 ºC. The cells were harvested at 5,000 rpm for 15 min and resuspended in 100 mM NaCl, 100 mM Tris pH 8.0 and disrupted through a microfluidizer. The lysate was then spun down at 20,000 rpm for 45 min and filtered. Protein was purified in two steps by Ni affinity chromatography and anion exchange chromatography using an ÄKTA system (GE Healthcare). The lysate was then loaded onto a 1 mL HisTrap HP column (GE Healthcare). The column was subsequently washed with 10% B and 20% B and eluted with 100% B. The Ni elution fraction was diluted 10-fold with 20 mM Tris pH 8.0 and was loaded onto a 1 mL HiTrap Q column (GE Healthcare). Elution was carried out by a 0-100% B gradient over 20 column volumes collecting 1.0 mL fractions. Flow rates were typically held constant at 1.0 mL/min or lowered if the pressure exceeded the limit of the column accordingly. HitrapQ fractions were further polished on gel filtration column superdex 200 16/60 in 20mM Tris pH 8.0, 150 mM NaCl. GST-tau was produced as previously reported (Min et al., 2010).

### HTS of p300 Inhibitors Based on the Homogeneous Time-Resolved Fluorescence Assay

50 nL of compound (final 0.5% DMSO) was added to 5 µL (final 6 nM) GST-tau in a 384-well plate. The reaction was initiated by adding 5 µL (final 1 nM) p300, followed by 1 h incubation at RT. At the end of the reaction, 10 µL/well of quench/detection mixture containing10 nM mAB359, 2.4 nM donor (anti-rabbit IgG-EuK), 3.6 nM acceptor (anti-GST-D2), and 25 µM anacardic acid (a known p300 inhibitor as the quench reagent) in detection buffer (50 mM sodium phosphate, pH7.9, 0.8 M KF) was added. The final mixture was then incubated at RT for another 2hr. After incubation, signal was read on EnVision Multilabel Plate Reader (PerkinElmer; ex: 340 nm, em: 665/620 nm). We used DMSO and anacardic acid as negative and positive controls, respectively. Percent inhibition was calculated as (FRET signal of DMSO-FRET signal of compound)/(FRET signal of DMSO-FRET signal of anarcardic acid) ×100%

To rule out compounds that suppress FRET through mechanisms other than inhibition of p300, we designed a counter screen in which compounds were added after 1 h incubation of the enzymatic reaction, followed immediately by the subsequent quench/detection. Any compounds that suppressed the FRET signal were considered false positives.

### Orthogonal MMBC Assay

The thiol-reactive dye MMBC ([10-(2,5-dihydro-2,5-dioxo-1H-pyrrol-1-yl)-9-methoxy-3-oxo-,methyl ester 3H-naphthol(2,1-b) pyran-S-carboxylic acid, known as ThioGlo 1] was used to detect CoA, the product of the acetylation reaction. Once MMBC reacts with CoA, it becomes fluorescent with excitation wavelength at 379 nm and emission wavelength at 513 nm. The assay was performed with a final volume of 50 μL in assay buffer (100 mM HEPES, pH7.5, 0.01% Triton-X100, 500 μM TCEP). Compounds of various concentrations were added to the mixture of 200 nM p300, 0.6 µM Tau, 100 µM Ac-CoA, and 250 µM MMBC. The reaction progresses were continuously monitored on plate reader Flexstation III (Molecular Devices) for 1 h. The slopes of the linear portion of the reaction were used to calculate the percentage of inhibition.

### Differential Scanning Fluorimetry

1 µM p300 with 5X syprorange was incubated with 25 µM, or 50 µM 37892 in the assay buffer. The Tm of p300 was monitored on real-time PCR machine Stratagene Mx3005p.

### HEK293T Cell Culture

HEK293T cells cultured in Dulbecco’s Modified Eagle’s medium (DMEM) with 10% fetal bovine serum were transfected with expression vectors encoding FLAG-tagged full-length hTau and myc-tagged p300, or empty vector, using lipofectamine 2000 (Life Technology). For mCherry-GFP-LC3 reporter, HEK293T cells were infected with lentivirus expressing mCherry-GFP-LC3, and single clone of infected cell with moderate expression was isolated to establish the reporter line.

### Primary Neuronal Cultures and Lentiviral or AAV Infections

Primary neuronal cultures were established from cortices of Sprague-Dawley rat pups (Charles River Laboratories) on postnatal day 0 or 1. Purified cells were plated at 600,000 cells/ml in Neurobasal medium supplemented with B27 (Invitrogen) on poly-ornithine-coated plates. All experiments were performed at 11-14 days in vitro (DIV) unless noted otherwise. Lentivirus was generated, purified, and used for infection as described (Chen et al., 2005). Briefly, recombinant lentivirus was produced by cotransfection of the shuttle vector (pRRL), two helper plasmids, delta8.9 packaging vector, and VSV-G envelope vector into HEK293T cells and purified by ultracentrifugation (Sun and Gan, 2011). AAV viral vectors expressing P301S tau were generated and purified from Viro-vek, Inc (Hayward, CA). All drug treatments and analyses were performed at DIV 10-13 days.

### Differentiation of Human iPSC–Derived Neurons

Human iPSC-derived neurons were differentiated with a simplified two-step protocol (pre-differentiation and maturation), as described previously (Wang et al. 2017). Briefly, iPSC line (WTC11) containing *Ngn2* transgene integrated to AAVS1 locus were pre-differentiated in Knockout Dulbecco’s modified Eagle’s medium (KO-DMEM)/F12 medium containing growth factors and doxycycline. After pre-differentiation, the neuronal precursor cells were matured in maturation medium containing 50% DMEM/F12, 50% Neurobasal-A medium with all the growth factors. Human neurons are assayed at 5-8 weeks post-differentiation.

### Tau secretion assay

Neuronal tau secretion assay was performed on matured neurons (DIV 11-14 for primary neuron, 5-8 weeks old for human iPSC-derived neurons). Fresh artificial cerebrospinal fluid (ACSF) containing NaCl (140 mM), KCl (5 mM), CaCl_2_(2.5 mM), MgCl_2_ (2 mM), HEPES (10 mM) and glucose (10 mM) was added to neurons to replace the original medium (neurobasal medium with B27 supplement). After 3h incubation at 37 °C, the conditioned medium was harvested and centrifuged at 10,000 RPM to remove cell debris, and subjected to hTau ELISA analysis to quantify the amount of the extracellular tau. For HEK293T cells, serum-free DMEM was used to replace post-transfection medium, and incubated at 37 °C for 24-48 h. The conditioned medium was harvested and centrifuged at 10,000 RPM to remove cell debris, and concentrated by 25-fold using Amicon Ultra-0.5 Centrifugal Filter Unit with Ultracel-30 membrane (Millipore). The concentrated medium was subjected to immunoblot analysis to quantify the amount of extracellular t-tau, p-tau and ac-tau. LDH assay (Promega) was used to monitor cell toxicity per manufacturer’s protocol.

### Homogenization of Cells and Tissues for Immunoblot Analyses

HEK293T cells or Mouse brain tissues were homogenized in RIPA buffer containing protease inhibitor cocktail (Sigma), 1 mM phenylmethyl sulfonyl fluoride, phosphatase inhibitor cocktail (Sigma), 5 mM nicotinamide (Sigma) and 1 μM trichostatic-A (Sigma). Neurons were homogenized in N-Per buffer (Thermo Fisher) containing all the inhibitors as above. Mouse brain tissues were sonicated after homogenization. Lysates were centrifuged at 14,000 RPM at 4 °C for 15 min. Supernatants were collected and protein concentrations were determined by the BCA assay (Thermo Fisher). Pellets containing the nuclei were washed with 1/3 volume of the lysis buffer, and used for histone extraction with HCl (0.24 N) overnight incubation. The same amount of proteins was resolved on a 4-12 % SDS-PAGE gel (Invitrogen), transferred to PVDF membrane (Bio-Rad), and probed with appropriate antibodies. Bands in immunoblots were visualized by enhanced chemiluminescence (Pierce) and quantified by densitometry and ImageJ software (NIH). Representative blots from same gel/membrane were shown and compared in the same figure. Samples from non-adjacent lanes were separated by a line.

### ELISA

AcH3K18 ELISA was performed using EpiQuik Global Acetyl Histone H3K18 Quantification Kit (Colorimetric) (EpiGentek), according to manufacturer’s protocol. CSF samples were concentrated by 10-folds using Amicon ultra centrifugal filters 3K (Millipore). Sensitive hTau ELISAs were adapted and modified according to previous reports (Barten et al., 2011, Meredith et al., 2013) and described previously (Wang et al. 2017). Briefly, mouse monoclonal antibody HT7 or BT2 were used for capture. The respective analytes were detected with alkaline phosphatase–conjugated mouse monoclonal antibodies Tau5 (BioLegend). Recombinant full length hTau (rPeptide) were used to generated standard curves for each assay. The CDP-Star substrate (Invitrogen) is used as chemiluminescent alkaline phosphatase substrate.

### Immunofluorescence Staining

Cells grown on coverslips were washed with phosphate-buffered saline (PBS) and fixed in fresh 4% paraformaldehyde, and permeabilized with 0.1% Triton X-100. Coverslips were washed in PBST (phosphate-buffered saline, 0.01% Triton X-100), and incubated for 1 h in blocking solution containing PBST and 5% normal goat serum. The cells were then incubated in blocking solution containing primary antibody overnight at 4 °C, followed by incubation with secondary antibody for 1 h. Primary antibodies include MC1 (1:500), MAP2 (1:800) and acH3K18 (1:1000). Secondary antibodies include fluorescein-labeled goat anti-mouse IgG and goat anti-rabbit IgG (1:500, Vector Laboratories). Hoechst (Thermo Fisher) was used to label the nuclei. Images were acquired by Leica microscope (DM5000 B) and analyzed by Micro-Manager software (UCSF, San Francisco, CA). For *p300*^*F/F*^*/CBP*^*F/*^ and *p300*^*F/+*^*/CBP*^*F/+*^ neurons, fully automated ArrayScan high-content system (Thermo) were used to acquire images and quantify MC1 signal and neuronal health parameters.

### RNA Isolation and Quantitative Real-Time PCR (qRT-PCR)

Total RNA was isolated from *p300*^*F/F*^*/CBP*^*F/F*^ primary neurons with the Direct-zol™ RNA MiniPrep kit (Zymo), and the remaining DNA was removed by incubation with RNase-free DNase (Zymo). Purified messenger RNA was then converted to complementary DNA by the TaqMan reverse transcription (RT) kit (Applied Biosystems). Quantitative RT-PCR was performed on the ABI 7900 HT sequence detector (Applied Biosystems) with SYBR Green PCR master mix (Applied Biosystems). The average value of three replicates of each sample was expressed as the threshold cycle (Ct), at which the fluorescence signal starts to increase rapidly. Then, the difference (ΔCt) between the Ct value for hTau and the Ct value for mouse GAPDH (ΔCt = Ct(hTau)-Ct(GAPDH)) was calculated for each sample. The relative levels of gene expression for each sample was determined by 2-ΔCt and expressed as the-fold change. The following primers were used for quantitative RT-PCR: hTau (forward, 5’-GTTGGGGGACAGGAAAGATCAG-3’; reverse, 5’-CCGGGAGCTCCCTCATC-3’), mouse GAPDH (forward, 5’-GGGAAGCCCATCACCATCTT-3’; reverse, 5’-GCCTTCTCCATGGTGGTGAA-3’).

### Tau Purification and Fibrilization

Myc-tagged repeat domain (K18) of P301L tau was expressed in Terrific broth media containing sodium chloride (500 mM) and a small-molecule chaperone, betaine (10 mM), to improve expression and minimize degradation. Expression was induced with 200 μM IPTG for 3.5 h at 30 °C. Tau was purified as described (Barghorn et al., 2005). Purified tau was dialyzed overnight at 4 °C into aggregation assay buffer (Dulbecco’s PBS pH 7.4, 2 mM MgCl_2_, 1 mM DTT). Aggregation of tau (_2_0 μM) was induced by the addition of a freshly prepared heparin sodium salt solution (Santa Cruz) at a final concentration of 88 μg/ml. Tau fibril samples were labeled with Alexa Fluor 647 NHS ester (Molecular Probes) at 1:40 molar ratio of dye:tau monomer for 1 h at room temperature. Samples were centrifuged at 100,000 g for 1 h at 4 °C to remove unreacted free dye. Pellets containing tau fibrils were resuspended in Dulbecco’s PBS, pH 7.4, 2 mM MgCl_2_. Tau fibril concentrations were quantified by SDS-PAGE with a set of tau monomer standards.

### In Vitro Fibril-Induced Tau Spreading Model

The in vitro fibril-induced tau spreading model was adapted and modified according to previous report (Guo and Lee, 2013). Briefly, primary neurons cultured on coverslips were infected with AAV-P301S hTau on DIV3, and synthetic tau fibrils (K18/P301L, 100 nM) were added to the medium on DIV5. Neurons without AAV infection (AAV-) and neurons treated with PBS instead of fibrils were included as negative controls. Neurons were incubated with the fibril for 7-10 days. To test the effect of compound treatment, drugs were added to the culture medium together with the fibril, and replenish once after 3 days. Immunofluorescent staining using MC1 antibody (1:500) was performed to detect tau aggregates. Briefly, neurons grown on coverslips were washed with fresh medium containing 0.01% trypsin, and fixed in fresh 4% PFA with 0.1% Triton-X, followed by regular immunofluorescent staining procedure.

### Immunohistochemistry and Image Analysis

Anesthetized mice were transcardially perfused with 0.9% saline. Mouse brains were removed and fixed in 4% paraformaldehyde for 48 h. Fixed brains were cryopreserved in 30% sucrose in PBS for at least 2 days, and coronal brain sections (30 μm) were obtained with sliding microtome. For immunostaining, floating brain sections were permeabilized and incubated in blocking solution (10% normal goat serum in 0.3% Triton X-100 TBST) at room temperature for 1 h. Sections were then incubated with primary antibodies, including p-p300 (S1834) (1:100), AT8 (1:500), CKiδ (1:5000), Chmp2B (1:600), lamp1 (1:1000), acH3K18 (1: 1000) and MC1 (1:500). After overnight incubation, the sections were incubated with secondary antibodies including Cy3-labeled donkey anti-rabbit IgG (1:500, Jackson ImmunoResearch) and fluorescein-labeled goat anti-mouse IgG (1:500, Vector Laboratories). Images were acquired by LSM880 confocal system (Zeiss) with Airyscan and a 20x or 63x oil-immersion objective lens, or Keyence BZ-X7000 microscope, and analyzed using ImageJ software (NIH). Experimenters quantifying immunoreactivity were blind to the mouse genotype and treatment conditions.

### Mice

Mice were housed in a pathogen-free barrier facility 165 with a 12-h light/dark cycle and ad libitum access to food and water. All animal procedures were carried out under guidelines approved by the Institutional Animal Care and Use Committee of the University of California, San Francisco. P301Stau transgenic mice (PS19), *Ep300*^*F/F*^ and *CBP*^*F/F*^ mice were purchased from Jackson Laboratory. *Ep300*^*F/F*^ mice were crossed with *CBP*^*F/F*^ mice to generate *Ep300*^*F/F*^*/CBP*^*F/F*^ mice. PS19 mice were crossed to *Ep300*^*F/F*^*/CBP*^*F/F*^ mice to generate *tauP301S/p300*^*F/+*^*/CBP*^*F/+*^ mice, which were further crossed with *Ep300*^*F/F*^*/CBP*^*F/F*^ mice to generate *tauP301S/Ep300*^*F/F*^*/CBP*^*F/F*^ mice. Mice were assigned into gender-and age-matched treatment groups in a randomized manner for all experiments. The sample size for each experiment was determined based on previous experiences with these models.

### Stereotaxic Injection

Mice were anesthetized with 2% isoflurane by inhalation for the duration of surgery, and secured on a stereotaxic frame (Kopf Instruments). Two-month-old PS19 mice and non-transgenic control mice were injected stereotaxically at a rate of 0.5 μl/min, with equal amounts (1×10^12^ genomic particles) of AAV1 expressing either Cre or GFP control (ViroTek), into the dentate gyrus of right hippocampus. 2 μL of synthetic tau fibrils (K18/PL) was injected into CA1 region of left hippocampus. The following coordinates were used for dentate gyrus (anterior-posterior −2.1, medial-lateral −1.7, dorsal-ventral −2.1) and CA1 (anterior-posterior −2.5, medial-lateral +2.0, dorsal-ventral −1.8). Control animals were injected with 2 μL of PBS instead of fibril.

### Statistics

Data were analyzed with GraphPad Prism 7 (GraphPad) or Stata. Differences between means were assessed with paired or unpaired Student’s *t* test, one-way or two-way analysis of variance, followed by post hoc testing of pairwise comparisons among genotypes (with Tukey’s or Dunnett’s correction for one-way ANOVA and Bonferroni correction for two-way ANOVA), or by mixed effects model, as indicated. Pearson’s correlation coefficients were used to quantify the linear relationship between two variables. Outliers are pre-established as data outside of mean ± 2SD. The level of significance was set as p < 0.05.

## Supplemental Text and Figures

**Figure S1.**
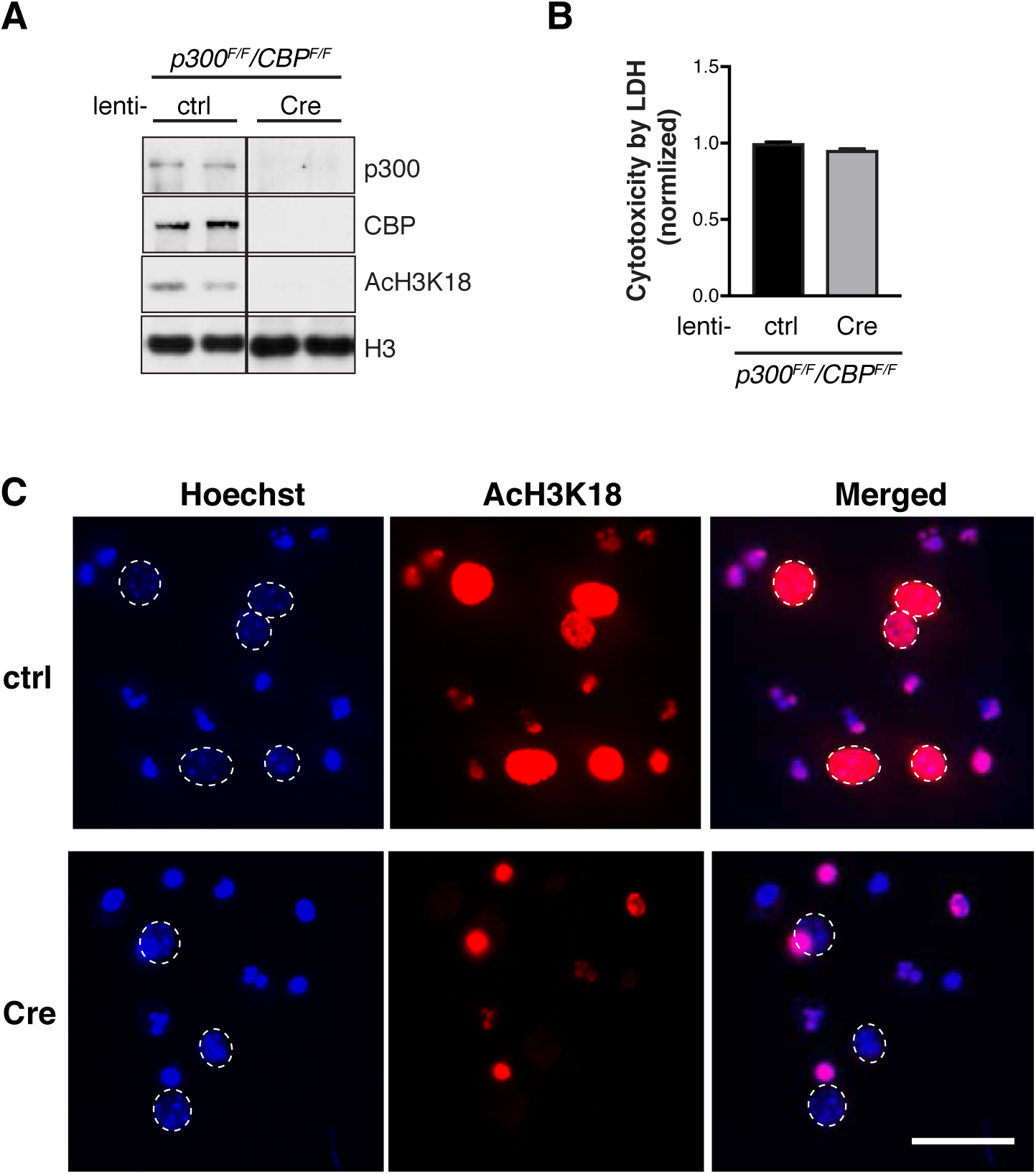
p300/CBP Double Knockout Abolishes AcH3K18 without Inducing Cytotoxicity in Neurons (Related to Figure 1, 2) (A) Representative immunoblots of p300, CBP, acH3K18, and H3 in lysates of p300^F/F^/CBP^F/F^ primary neurons infected with lenti-ctrl or lenti-Cre. (B) Cytotoxicity by LDH release assay, normalized to ctrl. n=4 wells from two independent experiments. (C) Representative immunofluorescence staining with anti-acH3K18 antibody and Hoechst in p300^F/F^/CBP^F/F^ primary neurons infected with lenti-ctrl or lenti-Cre. Scale bar: 20 μm.

**Figure S2.**
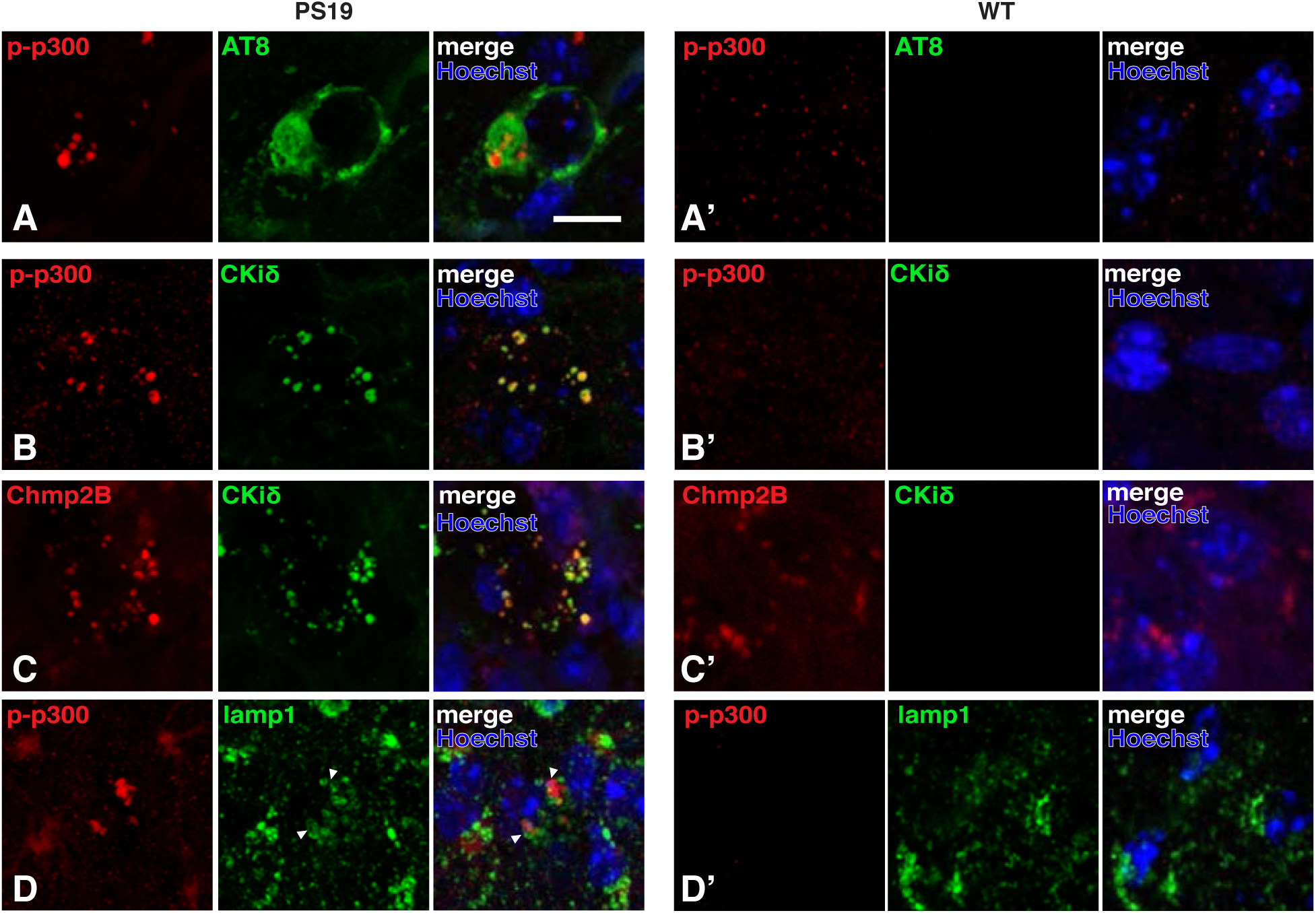
p-p300 (S1834) Accumulates in Granulovacuolar Lesions Resembling Autophagic Intermediates in Neurons with p-Tau Inclusion (Related to Figure 1) (A, B) Immunohistochemical staining of cortex of PS19 mice showing aberrant accumulation of (B)p-p300 (S1834) in neurons with AT8-positive tau inclusions (A), which is absent in WT mice (A′). (B) p-p300 signal co-localizes with casein kinase 1 (Ckiδ), a molecular marker for granulovacuolar degeneration body (GVB), which is absent in WT mice (B′). (C) Ckiδ-labeled GVB structures largely overlap with chmp2B (late-endosomal marker), suggesting p-p300 accumulates in the amphisome stage of autophagy (fusion of autophagosome and multivesicular body (MVB)). The co-localization is absent in WT mice (C′). (D) p-p300-containing granular structures (absent in WT mice, D′) are enveloped by lamp1 (lysosomal membrane marker), suggesting that p-p300 accumulates in autophagic intermediates with incomplete lysosome fusion. Scale bar: 10 μm.

**Figure S3.**
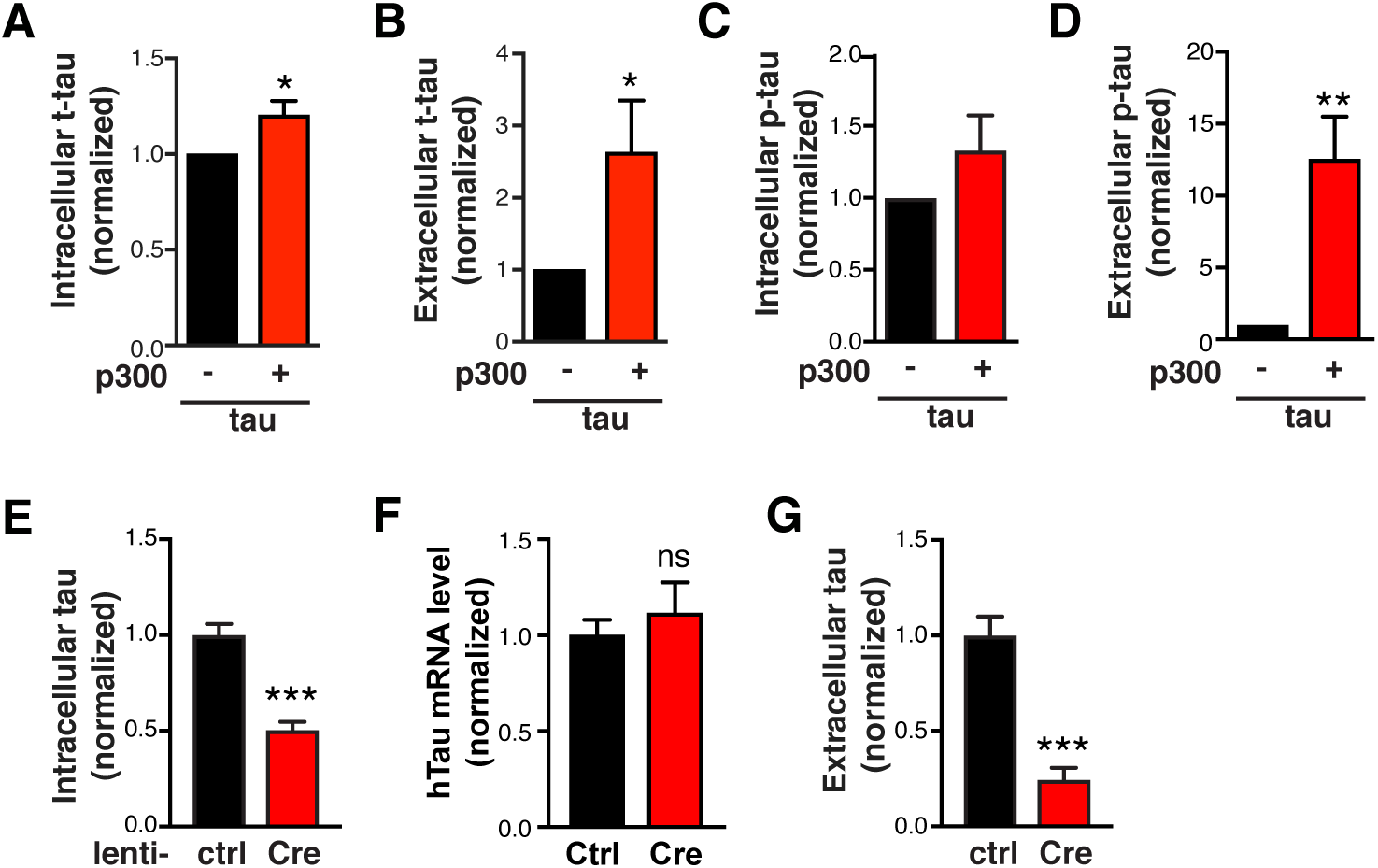
p300 Overexpression in HEK293T Cells Increases Tau Accumulation and Secretion, while p300/CBP Double-knockout in Primary Neurons Reduces Tau Accumulation and Secretion (Related to Figure 3) (A-D) Quantification of levels of t-tau (A, B) and p-tau (C, D) in the cell lysate (A, C) and conditioned medium (B, D) of HEK293T cells transfected with tau alone or tau+p300, normalized to control. n=8 wells from eight independent experiments. *p<0.05, **p<0.01 by unpaired t test. Values are mean ± SEM. (E, G) Tau levels in lysates and in conditioned medium (secreted over 3 h) of p300^F/F^/CBP^F/F^ primary neurons infected with lenti-ctrl or lenti-Cre. Tau levels were measured by ELISA and normalized to control. n=6 wells from three independent experiments. ***p<0.001 by unpaired t test. Values are mean ± SEM. (F)hTau mRNA levels in p300F/F/CBPF/F primary neurons infected with lenti-ctrl or lenti-Cre, by qRT-PCR. n=4 wells from two independent experiments. ns, non-significant by unpaired t test. Values are mean ± SEM.

**Figure S4.**
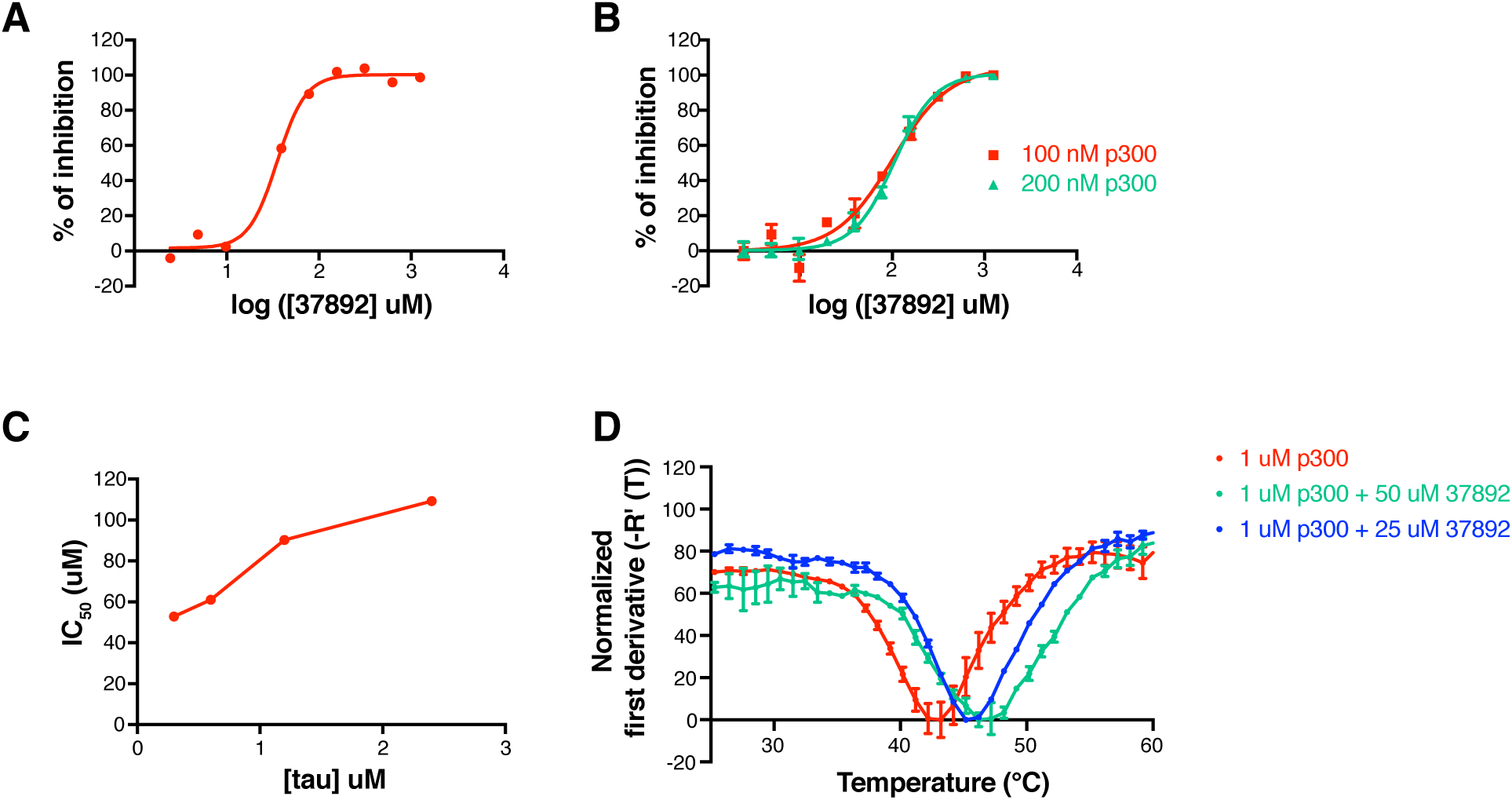
Characterization of 37892 as a p300 Inhibitor (Related to Figure 4) (A) 37892 inhibited tau acetylation by p300 with an IC_50_ of 35 μM. (B) Orthogonal MMBC assay confirms the inhibitory activity of 37892, albeit at a lower IC_50_ of 100 μM. IC_50_ of 37892 remained the same under different p300 concentrations, indicating that aggregation is not a likely mechanism of inhibition. (C) Dose-response curves of 37892 under different tau concentrations. (D)Differential scanning fluorimetry shows that 37892 binds to p300. The Tm of p300 increases from 42.5°C to 45.4°C and 47.2°C when 25 μM and 50 μM 37892 was added, respectively.

**Figure S5.**
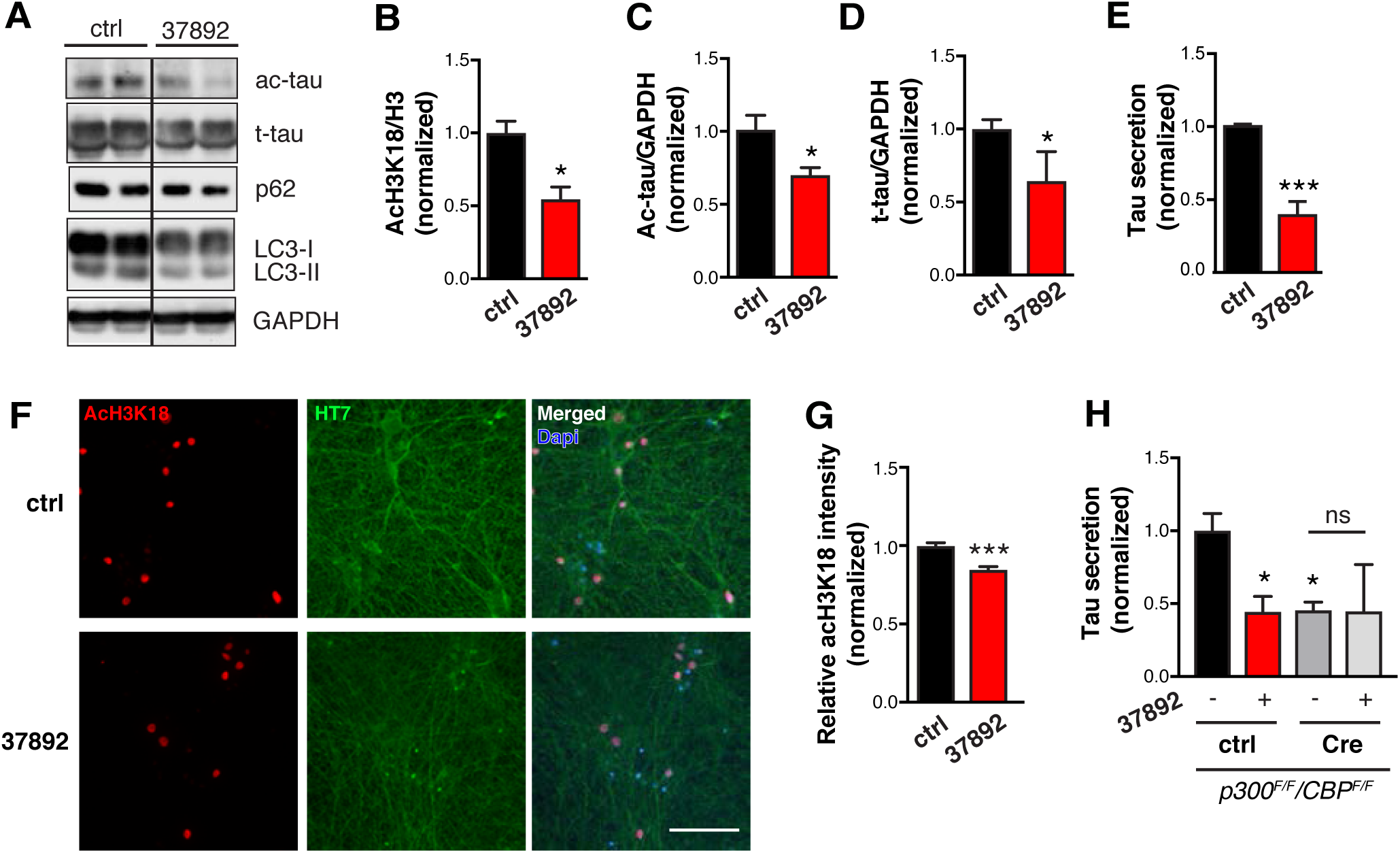
37892 Promotes Autophagic Flux and Reduces Tau Secretion in Human iPSC-induced Neurons (Related to Figure 4) (A) Representative immunoblot of acH3K18, H3, ac-tau (K174), t-tau (HT7), and GAPDH in lysates of human iPSC-induced neurons treated with DMSO (control) or 37892 (50 μM) for 3 days. Quantification of levels of acH3K18 (B) and ac-tau (C) and t-tau (D) after 3 days of treatment, normalized to ctrl. (E) Quantification of relative tau secretion over 3 h, based on levels of extracellular tau and intracellular tau measured by ELISA and normalized to control. n=6 wells from three independent experiments. *p<0.01, ***p<0.001 by unpaired t test. (F) Representative immunofluorescent staining with anti-AcH3K18 and HT7 antibody in human iPSC-derived neurons treated with ctrl (DMSO) or 37892 (50 μM). Scale bar: 100 μm. (G) Quantification of acH3K18 signal intensity relative to Dapi and normalized to control. n=64 cells (control) and 36 cells (37892). ***p<0.001 by unpaired t test. Values are mean ± SEM. (H) Quantification of relative tau secretion over 3 h in p300/CBP double knockout neurons treated with 37892. p300^F/F^/CBP^F/F^primary neurons were infected with lentivirus expressing empty vector (lenti-ctrl) or Cre recombinase (lenti-Cre), and treated with DMSO (control) or 37892 (50 μM) for 24 h. n=4 wells from 2 independent experiments. *p<0.05, ns, non-significant, by one-way ANOVA and Sidak’s multiple comparisons test. Values are mean ± SEM.

**Figure S6.**
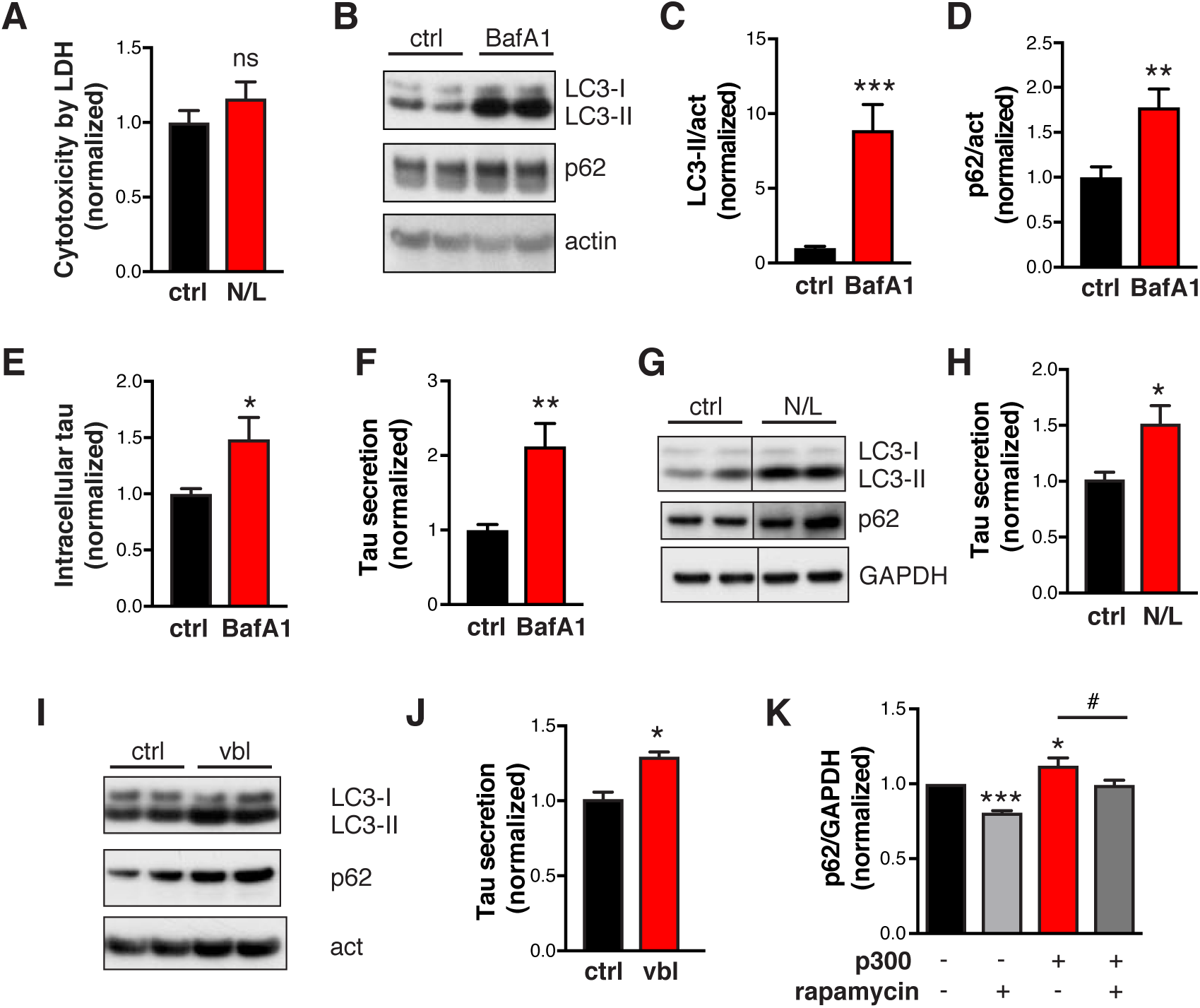
Primary Neurons and Human iPSC-derived Neurons Treated with N/L or BafA1, and HEK293T cells treated with rapamycin (Related to Figure 5, 6) (A) Quantification of cytotoxicity in hTau-expressing primary neurons treated with N/L using LDH release assay, normalized to ctrl. n=4 wells from 2 independent experiments. ns, non-significant by unpaired t test. (B–F) hTau-expressing primary neurons treated with BafA1. (B) Representative immunoblot of LC3-I, LC3-II, p62, and actin in lysates of rat primary neurons infected with AAV-P301S hTau, after 24 h of treatment with DMSO (ctrl) or BafA1 (10 nM). (C, D) Levels of LC3-II (C) and p62 (D) relative to actin and normalized to control. (E, F) Levels of intracellular tau (E) and relative tau secretion over 3 h (F), measured by ELISA and normalized to control. n=6 wells from three independent experiments. *p<0.05, **p<0.01, ***p<0.001 by unpaired t test. Values are mean ± SEM. (G-J) N/L and vinblastine (vbl) block autophagic flux and increases tau secretion in human-iPSC-derived neurons. (G, I) Representative immunoblot of LC3-I, LC3-II, p62, and GAPDH or actin in lysates of 8–10-week-old human neurons, after 24 h of treatment with DMSO (ctrl), N/L (20 mM NH_4_Cl and 200 μM leupeptin), or vinblastine (vbl, 5 μM). (H, J) Quantification of relative tau secretion with N/L or vbl treatment, based on levels of extracellular tau and intracellular tau measured by ELISA and normalized to control. n=4 wells from two independent experiments. *p<0.05, unpaired t test. Values are mean ± SEM. (K) Quantification of p62 levels in lysates of HEK293T cells transfected with tau alone or tau + p300 and treated with DMSO (ctrl) or rapamycin (1 μM) for 24 h. *, #p<0.05, ***p<0.001 by one-way ANOVA and Sidak’s multiple comparisons test. Values are mean ± SEM.

**Figure S7.**
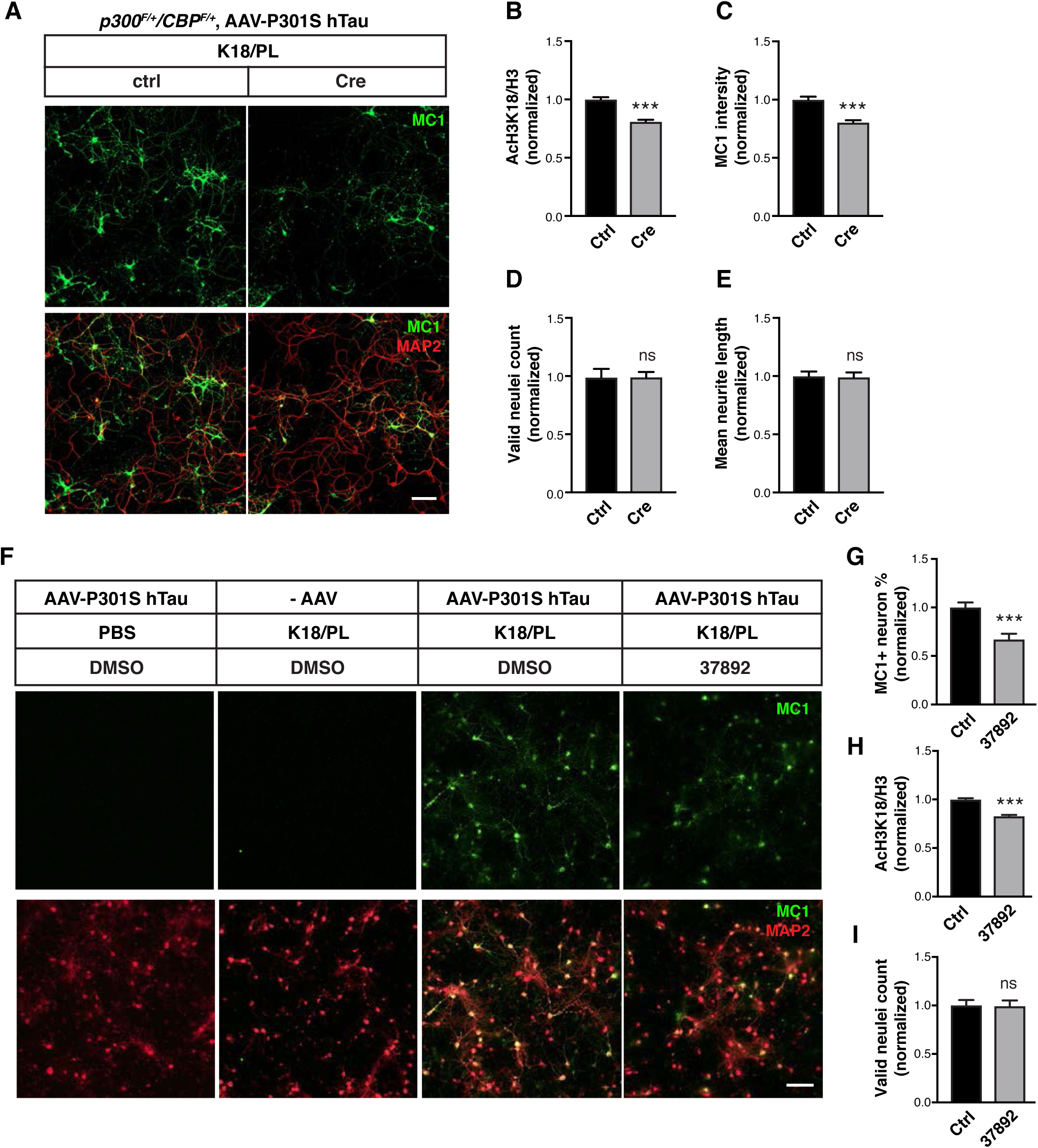
p300/CBP Heterozygous Knockout and 37892 Treatment Reduce Fibril-Induced Tau Spreading in Primary Neurons (Related to Figure 7) (A–E) p300/CBP heterozygous knockout reduces fibril-induced tau spreading in primary neurons. (A) Representative immunofluorescence staining with MC1 and MAP2 antibody in p300F/+ CBPF/+ primary neurons infected with AAV-P301S hTau and lenti-control or lenti-Cre, and treated with synthetic tau fibrils (K18/PL, 100 nM). Scale bar: 100 μm. (B–E) Quantification of acH3K18 signal (B), MC1 intensity (C), number of valid (live) nuclei (D), and mean neurite length (E), normalized to control. ***p<0.001, unpaired t test. n=12 wells from two independent experiments. Values are mean ± SEM. (F–I) 37892 treatment reduces fibril-induced tau spreading in primary neurons. (F) Representative immunofluorescence staining with MC1 and MAP2 antibody in primary rat neurons infected with AAV-P301S hTau and treated with synthetic tau fibril (K18/PL, 100 nM) and 37892 (50 μM) or DMSO (control). Negative controls (PBS-treated, AAV non-infected) are included. Scale bar: 100 μm. (G) Quantification of percentage of MC1-positive neurons, normalized to control. (H) Quantification of acH3K18 signal intensity relative to Hoechst and normalized to control. n>800 cells/treatment. (I) Number of valid (live) nuclei, normalized to control. n=13 fields from two independent experiments. ***p<0.001, unpaired t test. Values are mean ± SEM.

